# Nuclear quality control of non-imported secretory proteins attenuates proteostasis decline in the cytosol

**DOI:** 10.1101/2023.06.27.546668

**Authors:** Papiya Banik, Janine Kamps, Qi-Yin Chen, Hendrik Luesch, Konstanze F. Winklhofer, Jörg Tatzelt

## Abstract

Mistargeting of secretory proteins to the cytosol can induce formation of aggregation-prone conformers and subsequent proteostasis decline. We have identified a quality control pathway that redirects non-ER-imported prion protein (PrP) to proteasomal degradation in the nucleus to prevent formation of toxic aggregates in the cytosol. Upon aborted ER import, PrP sequentially interacted with VCP/p97 and importins, which kept PrP soluble and promoted its nuclear import. In the nucleus, RNA buffered aggregation of PrP to facilitate ubiquitin-dependent proteasomal degradation. Notably, the cytosolic interaction of PrP with VCP/p97 and its nuclear import were independent of ubiquitination but required the intrinsically unstructured N-terminal domain of PrP. Transient proteotoxic stress promoted the formation of self-perpetuating PrP aggregates in the cytosol, which disrupted further nuclear targeting of PrP and compromised cellular proteostasis. Our study delineates a VCP/p97-dependent nucleus-based quality control pathway of non-ER-imported secretory proteins and emphasizes the important role of the nuclear milieu for the degradation of aggregation-prone proteins.

## INTRODUCTION

To maintain cellular protein homeostasis and to preclude toxic effects of aberrant protein conformers, components of the proteostasis network ensure proper protein folding as well as recognition and degradation of misfolded and non-functional proteins. The accumulation of protein aggregates in various neurodegenerative diseases suggests that an overload of quality control pathways and/or a decline in their efficiencies play a crucial role in the pathogenesis of these diseases (Hartl, 2017, Mogk et al., 2018, Schopf et al., 2017, Taipale et al., 2010). Another determinant of protein folding and aggregation is the chemical milieu of the cellular compartments. For example, the folding of membrane proteins and secretory proteins, which often contain disulfide bonds, is critically dependent on the specialized environment in the endoplasmic reticulum (Braakman & Hebert, 2013, Brodsky & Skach, 2011, Jessop et al., 2004). As a consequence, mislocalization of these proteins to the cytosol can trigger their misfolding and lead to the formation of toxic protein aggregates (Hegde & Zavodszky, 2019, Juszkiewicz & Hegde, 2018).

A prominent example of a secretory protein with neurotoxic potential is the mammalian prion protein (PrP). Prion diseases are invariably fatal neurodegenerative disorders in humans and animals caused by the conformational transition of the cellular PrP (PrP^C^) into pathogenic conformers, denoted scrapie prion protein (PrP^Sc^) that are infectious and neurotoxic (Aguzzi & Polymenidou, 2004, Prusiner, 1998, Wadsworth & Collinge, 2011). PrP^C^ is a classical secretory protein with a cleavable N-terminal signal peptide. After import into the ER, the C-terminus is modified with a glycosylphosphatidylinositol (GPI) anchor, which targets PrP^C^ to the outer leaflet of the plasma membrane (Stahl et al., 1987). However, several studies *in vitro*, *in cellulo*, in transgenic mice and patients suffering from prion diseases revealed that inefficient or altered translocation of PrP into the ER lumen can lead to the formation of PrP conformers that are neurotoxic but not infectious. From these studies the concept emerged that PrP-mediated neurodegeneration and propagation of infectious prions are linked to distinct pathogenic PrP conformers (Benilova et al., 2020, Chakrabarti et al., 2009, Chiesa et al., 2003, Hegde et al., 1999, Sandberg et al., 2011, Sandberg et al., 2014).

Transgenic mice expressing PrP devoid of the N-terminal ER-targeting signal developed severe ataxia and cerebellar degeneration and provided the first evidence that cytosolically localized PrP (cytoPrP) has a neurotoxic activity (Ma et al., 2002). Subsequently, toxic effects of mislocalized PrP in the cytosol were corroborated in different transgenic mice and cell culture models (Chakrabarti & Hegde, 2009, Deriziotis et al., 2011, Heller et al., 2003, Kristiansen et al., 2007, Kristiansen et al., 2005, Ma et al., 2002, Norstrom et al., 2007, Rambold et al., 2006, Rane et al., 2004). Moreover, intraneuronal accumulation of PrP^Sc^ is also described in some clinicopathological phenotypes of sporadic Creutzfeldt-Jakob disease (sCJD) (Parchi et al., 2012). Notably, wildtype PrP^C^ also forms cytosolic aggregates after transient proteasomal inhibition (Drisaldi et al., 2003, Ma & Lindquist, 2001, Ma & Lindquist, 2002, Yedidia et al., 2001). Initially, it was assumed that PrP is subjected to the endoplasmic reticulum (ER)-associated degradation (ERAD) pathway. This pathway involves recognition of non-native polypeptides in the ER lumen by ER-resident proteins and retrograde transport to the cytosol for proteasomal degradation (rev. in (Ellgaard & Helenius, 2003, Meusser et al., 2005, Nakatsukasa & Brodsky, 2008). However, it turned out that cytoPrP is generated from a fraction of PrP that is not imported into the ER, due to its inefficient ER signal peptide and the extended intrinsically disordered N-terminal domain (Drisaldi et al., 2003, Hegde & Bernstein, 2006, Heller et al., 2003, Heske et al., 2004, Kim et al., 2001, Miesbauer et al., 2009, Rambold et al., 2006, Rane et al., 2004, Rutkowski et al., 2001). Whereas the mechanisms associated with the ERAD pathway have been studied in great detail, quality control pathways operating specifically for non-imported secretory proteins are not well understood. In this study we have identified a VCP/p97-dependent quality control pathway that targets mislocalized PrP to the nucleus for proteasomal degradation and thereby prevents the accumulation of toxic PrP aggregates in the cytosol.

## RESULTS

### Non-ER-imported PrP is targeted to the nucleus for proteasomal degradation

To specifically study cytosolic quality control pathways that operate on the fraction of secretory proteins that were not translocated through the Sec61 translocon into the ER, we followed up four independent approaches to generate non-imported prion protein (Fig. 1A). 1. N3PrP has a mutation in its signal sequence that prevents binding of the nascent chain to the SRP and/or gating and thereby ER import (Kim et al., 2002). 2. Intrinsically disordered domains require alpha-helical domains in addition to the N-terminal signal peptide for efficient Sec61-mediated transport into the ER. As a consequence, ER import of a PrP variant composed only of the intrinsically disordered N-terminal domain of PrP (PrP115X) is strongly impaired (Dirndorfer et al., 2013, Gonsberg et al., 2017, Heske et al., 2004, Miesbauer et al., 2009, Miesbauer et al., 2010, Mohammadi et al., 2020, Pfeiffer et al., 2013, Zanusso et al., 1999). 3. Cyclic depsipeptide natural products of the Apratoxin class reversibly inhibit cotranslational translocation of secretory proteins through the Sec61 translocon, with Apratoxin S9 being the most potent synthetic analogue to date (Chen et al., 2014, Itskanov et al., 2023, Liu et al., 2009). 4. Thapsigargin, a non-competitive inhibitor of the sarco/endoplasmic reticulum Ca^2+^ ATPase, induces ER stress that activates a preemptive quality control pathway to reduce the import of secretory proteins into the ER (Hegde & Kang, 2008, Kang et al., 2006, Oyadomari et al., 2006). First, we studied N3PrP, a full-length PrP variant equipped with an N-terminal ER signal peptide and a C-terminal GPI signal sequence that is not imported into the ER after synthesis because of a mutation in its ER signal peptide (Kim et al., 2002). Interestingly, N3PrP was not only present in the cytosol of human neuroblastoma SH-SY5Y cells but also in the nucleus. Moreover, after treating the cells with the reversible proteasomal inhibitor Bortezomib, nuclear-localized N3PrP was strongly stabilized (Fig. 1B). Next, we analyzed the fate of non-imported PrP115X. After SRP-mediated targeting to the Sec61 translocon, a significant fraction of this PrP mutant is not translocated into the ER lumen and instead released into the cytosol. Immunocytochemistry followed by fluorescence microscopy of SH-SY5Y cells transiently expressing PrP115X showed only a weak signal, consistent with previous results indicating that non-imported PrP115X is rapidly degraded by the proteasome (Fig. 1C). Strikingly, upon Bortezomib treatment, PrP115X predominantly accumulated in the nucleus, revealing that non-ER-imported PrP was targeted to the nucleus for proteasomal degradation (Fig. 1C). To study the fate of wildtype PrP under conditions of blocked ER import, we expressed PrP lacking the C-terminal GPI-anchor signal sequence (PrPΔGPI). The ER import of PrPΔGPI is comparable to that of GPI-anchored PrP, but PrPΔGPI is secreted (Rambold et al., 2006, Winklhofer et al., 2003). Thus, by using this mutant we can make sure that cytosolic localization of PrPΔGPI is a consequence of its impaired ER import and not of PrP misfolding in the ER lumen (Satpute-Krishnan et al., 2014), nor endocytosis of PrP from the plasma membrane (Sunyach et al., 2003). Fluorescence microscopy suggested the localization of PrPΔGPI within the secretory pathway of control cells, as expected for a secreted protein. After inhibiting Sec61-mediated ER import by Apratoxin S9 for 4 h, PrPΔGPI was mostly detected in the nucleus (Fig. 1D). To facilitate the microscopic analysis, we equipped PrPΔGPI with a C-terminal GFP and generated mouse neuroblastoma N2a cell lines stably expressing PrPΔGPI-GFP. The activity of Apratoxin S9 to potently inhibit ER import was validated by immunoblotting of cell lysates (Fig. 1E). The upper band, representing N-linked glycosylated PrPΔGPI-GFP within the secretory pathway, was completely absent in cells exposed to Apratoxin S9 for 4 h. In addition, the amount of PrPΔGPI-GFP in conditioned media was significantly decreased (Fig. 1E). Similarly to untagged PrPΔGPI, after inhibiting ER import, PrPΔGPI-GFP was transported from the cytosol to the nucleus (Fig. 1F). To verify that the transport of non-ER-imported PrP from the cytosol into the nucleus is not a specific feature of established cell lines, we analyzed mouse primary cortical neurons transiently transfected with PrPΔGPI-GFP and treated with Apratoxin S9 by fluorescence microscopy. In line with our findings in established cell lines, non-ER-imported PrP was targeted to the nucleus of primary neurons (Fig. 1G). Previous work revealed that ER stress results in an overall reduced rate of ER import of secretory proteins (Hegde & Kang, 2008, Kang et al., 2006, Oyadomari et al., 2006). Specifically, during acute ER stress PrP has been shown to be co-translocationally targeted to proteasomal degradation (Kang et al., 2006, Orsi et al., 2006). However, in these studies the stabilization of PrP after proteasomal inhibition was only shown by Western blotting without addressing the cellular compartment of PrP accumulation. To assess the cellular localization of non-translocated PrP upon ER stress, we analyzed transiently transfected cells treated for 4 h with Thapsigargin. Fluorescence microscopy revealed that during ER stress a fraction of PrPΔGPI-GFP was not imported into the ER and targeted to the nucleus (Fig. 1H). While the four different approaches described above revealed that non-ER-imported PrP is targeted to the nucleus for proteasomal degradation, the relative amounts of non-translocated PrP were increased either by employing PrP mutants or during stress conditions. Notably, even under physiological conditions around 10%–20% of PrP^C^ are not imported into the ER and mislocalize to the cytosol (Rane et al., 2004). Thus, we were wondering whether this fraction is also targeted to the nucleus. To this end we analyzed the cellular localization of PrPΔGPI-GFP in stably expressing N2a cells up to 72 h after plating. After 24 h, PrP was not detected in the nucleus. However, after 48 h some cells showed a GFP signal in the nucleus, and by 72 h many nuclei contained PrPΔGPI-GFP, indicative of nuclear targeting of non-ER-imported PrP. We could not observe that nuclear localization of PrP decreased cellular viability (Fig. EV1). In sum, our five-pronged approach revealed that non-ER-imported PrP, which accumulates as a consequence of inefficient or impaired ER import, is targeted to the nucleus for proteasomal degradation.

**Figure 1.**
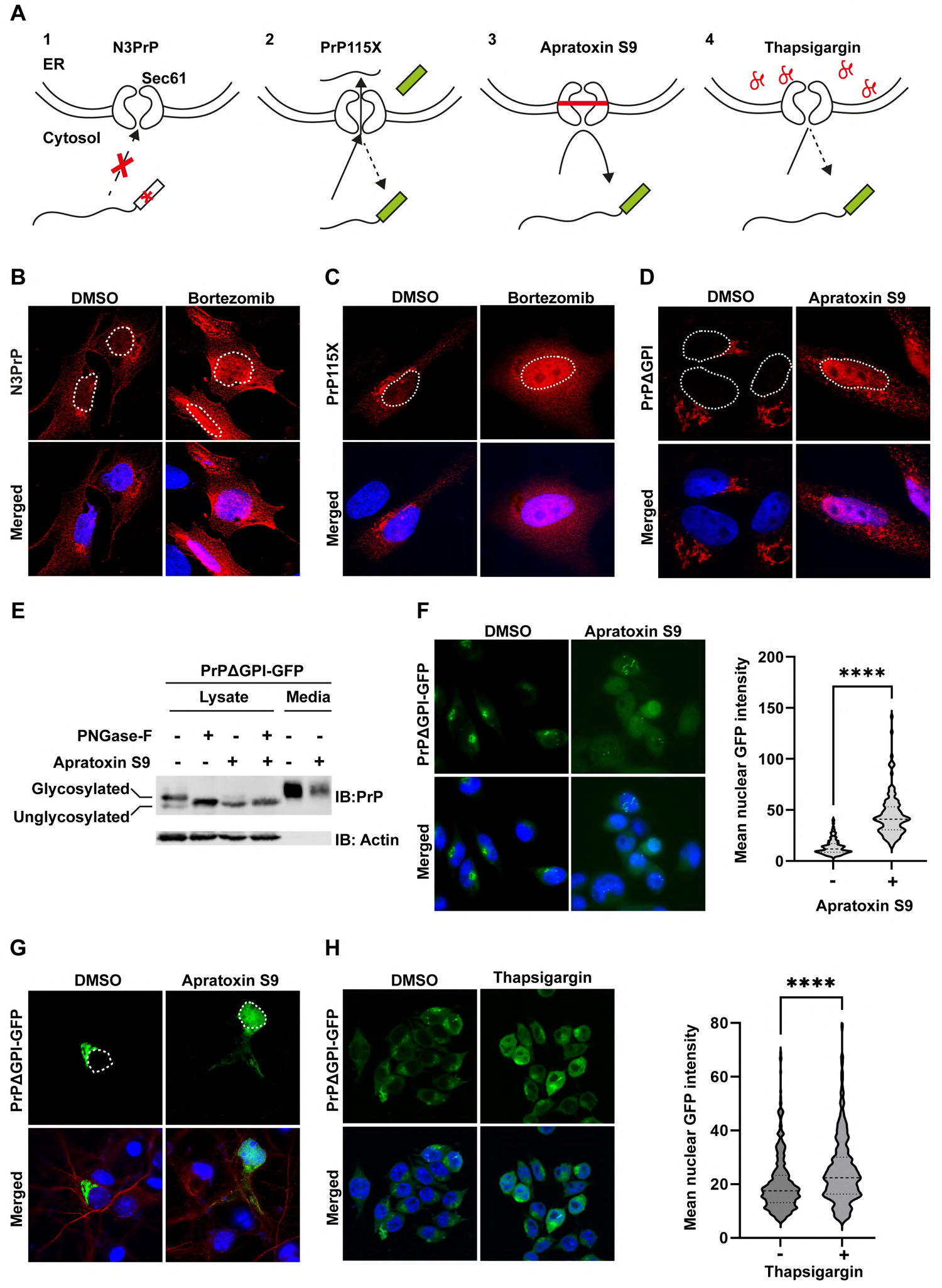
Non-imported prion protein is targeted to the nucleus for proteasomal degradation. **A. Scheme of the experimental approaches to generate non-imported PrP.** 1. N3PrP has a mutation in its signal peptide preventing its interaction with the signal recognition particle ER. 2. Deletion of the C-terminal structured domain impairs ER import of PrP115X. 3. Apratoxin S9 binds to and prevents translocation through the Sec61 translocon. 4. ER stress, induced by Thapsigargin, activates a preemptive quality control pathway that reduce import of secretory proteins into the ER. **B, C, D, E, F, G, H. Non-imported PrP is targeted to the nucleus in different cell lines and under different conditions.** **B, C,** SH-SY5Y cells transiently expressing N3PrP ((B) or PrP115X (C) were treated with DMSO (control), or the proteasome inhibitor Bortezomib (1µM) for 3 h and fixed, stained with antibodies against PrP and analyzed by SR-SIM. White dotted lines indicate boundaries of the nuclei. Nuclei were stained with DAPI (Merged). **D.** SH-SY5Y cells transiently expressing PrPΔGPI were treated with Apratoxin S9 (100 nM) for 4 h and fixed, stained with antibodies against PrP and analyzed by SR-SIM. White dotted lines indicate boundaries of the nuclei. Nuclei were stained with DAPI (Merged). **E.** HEK-293T cells were transiently transfected with PrPΔGPI-GFP. 24 h post-transfection, cells were treated for 4 h with either DMSO or Apratoxin S9 (100 nM) in fresh media. Cells were lysed and digested with PNGase-F or left untreated. Cell lysates and conditioned media were analyzed by immunoblotting using antibodies against PrP and β-Actin (loading control). Glycosylated and unglycosylated PrP species are indicated. **F.** N2a cells stably expressing PrPΔGPI-GFP were plated and treated for 4 h with either DMSO or Apratoxin S9 (100 nM), fixed and GFP fluorescence was analyzed by SR-SIM (left panel). Nuclei were stained with DAPI (Merged). Using FIJI software, the mean nuclear intensity of GFP fluorescence was quantified. The indicated line is the mean of the data set and was analyzed by two-tailed Mann Whitney test at 99% confidence interval, *** p=0.0001. At Least 50 cells were analyzed per biological replicate (n=3) (right panel). **G.** Primary cortical mouse neurons at DIV 6 were transfected with PrPΔGPI-GFP. 48 h post transfection the cells were treated for 4 h with either DMSO or Apratoxin S9 (100 nM), fixed, stained with antibodies against MAP2 and analyzed by SR-SIM. White dotted lines indicate boundaries of the nuclei. Nuclei were stained with DAPI (Merged). **H.** N2a cells stably expressing PrPΔGPI-GFP were treated for 4 h with either DMSO or Thapsigargin (5 µM) to induce ER stress. The cells were fixed and GFP fluorescence was analyzed by SR-SIM (left panel). Nuclei were stained with DAPI (Merged). Using FIJI software, the mean nuclear intensity of GFP fluorescence was quantified. The indicated line is the mean of the data set and was analyzed by two-tailed Mann Whitney test at 99% confidence interval, *** p=0.0001, **** p=0.00001. At least 50 cells were analyzed per biological replicate (n=3) (right panel).

### VCP/p97 and importin-ß are required for nuclear import of cytosolically localized prion protein

The preceding findings raised the question of the mechanism underlying the transport of PrP into the nucleus. In the ERAD pathway VCP/p97 is required for the retrograde translocation of misfolded secretory proteins from the ER and their targeting to cytosolic proteasomes (Neuber et al., 2005, Schuberth & Buchberger, 2005, Ye et al., 2003). To study whether VCP/p97 is involved in nuclear targeting of non-ER-imported PrP, we treated cells transiently expressing PrPΔGPI-GFP with Apratoxin S9 and VCP/p97 inhibitors. Indeed, inhibiting VCP/p97 by three different inhibitors, DBeQ, CB-5083 or NMS-873, decreased the nuclear localization of PrP and increased the formation of cytosolic PrP aggregates (Fig. 2A). A quantitative analysis using the stably PrPΔGPI-GFP-expressing cell line verified that nuclear targeting of non-ER-imported PrP in DBeQ-treated cells was significantly reduced (Fig. 2B). To analyze the role of VCP/p97 in more detail, we immunoprecipitated VCP/p97 under non-denaturing conditions to maintain protein-protein interaction and then analyzed co-purifying proteins for the presence of PrPΔGPI by Western botting. Since the interaction of VCP/p97 with its client proteins is only transient, we included TrapVCP in our experiments, which is a VCP/p97 mutant (E578Q) that does not release clients after binding (Hulsmann et al., 2018). Immunoblots of the input lysates confirmed that Apratoxin S9 efficiently blocked ER import. The upper band of PrPΔGPI, representing glycosylated species in the secretory pathway, was absent in lysates prepared from Apratoxin S9-treated cells (Fig. 2C). An interaction of PrPΔGPI with wildtype VCP/p97 could not be detected by this approach, neither in control nor in Apratoxin S9-treated cells. However, PrPΔGPI co-purified with TrapVCP when the cells had been treated with Apratoxin S9 to block ER import (Fig. 2C). Of note, TrapVCP did not interact with PrP in control cells without Apratoxin S9, revealing that the interaction was specific for non-ER-imported PrP. To address the question whether PrPΔGPI interacts with VCP/p97 only in the cytosol or also in the nucleus, we targeted PrP-GFP either to the nucleus or the cytosol by exchanging the ER signal peptide with a nuclear localization (NLS) or nuclear export signal (NES). Fluorescence microscopy confirmed that NES-PrP-GFP and NLS-PrP-GFP are almost exclusively located in the cytosol and nucleus, respectively (Fig. 4C). Although TrapVCP was present in both the nucleus and the cytosol (Fig. EV2), the co-immunoprecipitation experiments indicated that TrapVCP only interacted with NES-PrP-GFP but not with NLS-PrP-GFP (Fig. 2D).

**Figure 2.**
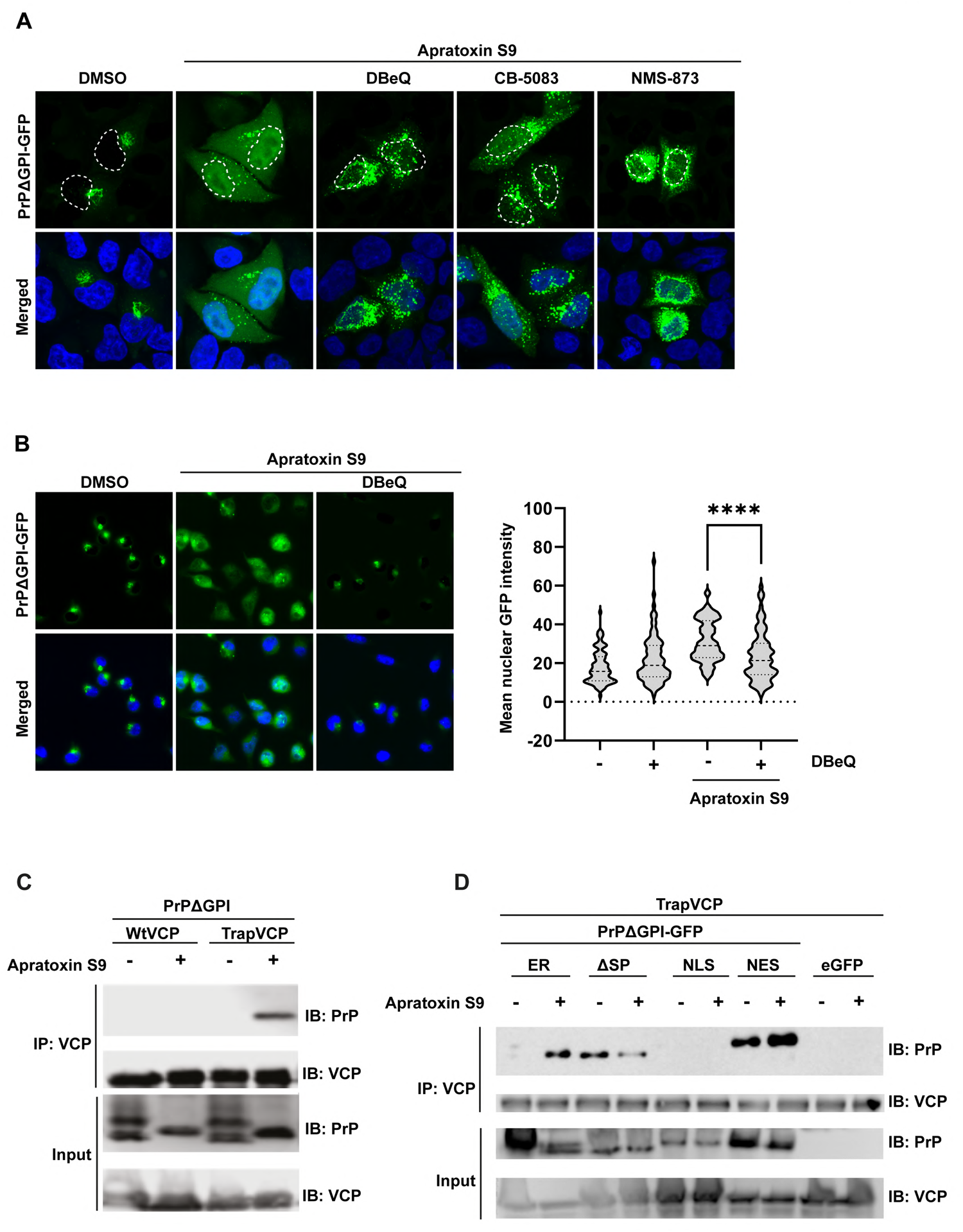
Inhibition of VCP/p97 interferes with nuclear targeting of non-imported PrP. **A, B. Pharmacological inhibition of VCP/p97 decreases nuclear import and increases cytosolic accumulation of non-imported PrP.** HeLa cells transiently transfected with PrPΔGPI-GFP were treated for 3 h with Apratoxin S9 (100 nM), or with Apratoxin S9 together with the VCP/p97 inhibitors DBeQ (10µM), or NMS-873 (10 µM) or CB-5083 (10 µM). Please note that the VCP inhibitors were added 1 h before Apratoxin S9. The cells were fixed and GFP fluorescence was analyzed by SR-SIM. Nuclei were stained with DAPI (Merged). White dotted lines indicate boundaries of the nuclei. **B.** N2a cells stably expressing PrPΔGPI-GFP were treated for 3 h with Apratoxin S9 (100nM), or with Apratoxin S9 together with DBeQ (10 µM). Please note that DBeQ was added to the cell culture media 1 h before Apratoxin S9. The cells were fixed and GFP fluorescence was analyzed by SR-SIM. Nuclei were stained with DAPI (Merged). Using FIJI software, the mean nuclear intensity of GFP fluorescence was quantified. The indicated line is the mean of the data set and was analyzed by Kruskal-Wallis test followed by Dunn’s Multiple Comparison Test at 99% confidence interval, *** p=0.0001, **** p=0.00001. At Least 50 cells were analyzed per biological replicate of n=3. **C, D. VCP interacts with cytosolic but not nuclear PrP.** **(C)** HEK-293T cells were co-transfected with PrPΔGPI and Strep-tagged WtVCP or E578QVCP (TrapVCP). 24 h post-transfection, cells were treated with DMSO or Apratoxin S9 (100 nM) for 4 h, lysed and subjected to immunoprecipitation under native conditions with Streptactin magnetic beads. The immunoblots were tested for antibodies against PrP (3F4) and VCP. **D.** HEK-293T cells were co-transfected with Strep-tagged TrapVCP and the PrP constructs indicated. 24 h post-transfection, cells were treated with DMSO or Apratoxin S9 (100 nM) for 4 h, lysed and subjected to immunoprecipitation under native conditions using Streptactin magnetic beads. Precipitated proteins were then detected by immunoblotting using antibodies against PrP and VCP/p97.

While VCP/p97 was obviously required for nuclear targeting of non-ER-imported PrP, we wondered whether importins play a role in this context. Cells transiently expressing PrPΔGPI were treated with Apratoxin S9 in combination with different importin inhibitors. In cells treated with Importazole, an inhibitor of ß-importins, non-ER-imported PrPΔGPI accumulated in the cytosol (Fig. 3A). Ivermectin, which inhibits α-importins, had no obvious effect on the nuclear translocation of PrP (Fig. 3A). A quantitative analysis in cells stably expressing PrPΔGPI-GFP confirmed that in the presence of Importazole the nuclear import of PrPΔGPI-GFP in Apratoxin S9-treated cells was significantly impaired (Fig. 3B). Taken together, these experiments revealed that after an aborted ER import efficient targeting of PrP to the nucleus is dependent on both VCP/p97 and ß-importins.

**Figure 3.**
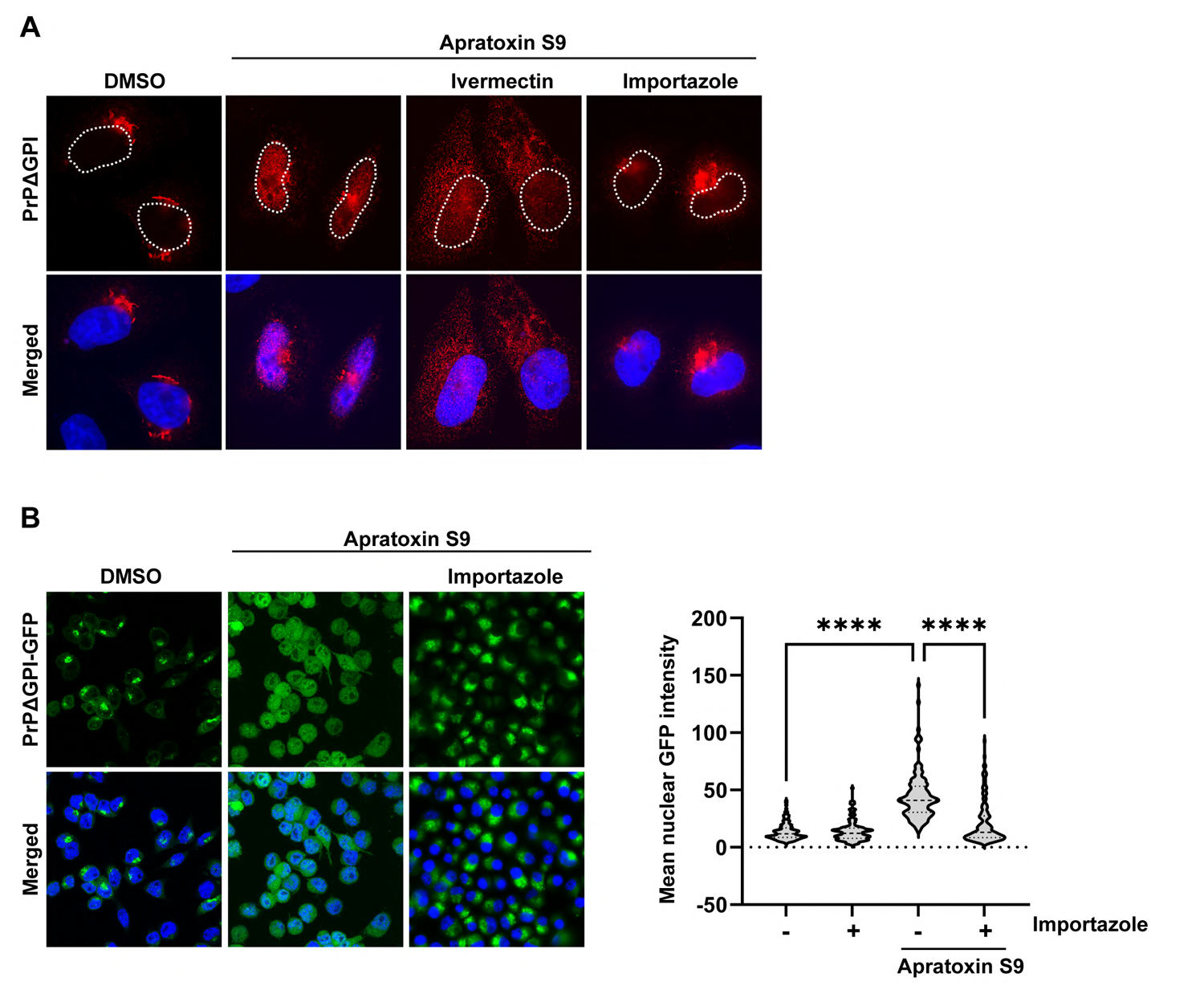
Nuclear import of PrP is importin-dependent. **A, B. Importin-β mediates nuclear import of PrP.** **(A)** HeLa cells were transiently transfected with PrPΔGPI. 24 h post-transfection cells were treated for 3 h with Apratoxin S9 (100 nM), or Apratoxin S9 with Ivermectin (50 µM) or Importazole (50 µM). Note that importin inhibitors were added 1 h before Apratoxin S9. Cells were fixed, stained with antibodies against PrP and analyzed by SR-SIM. Nuclei were stained with DAPI (Merged). White dotted lines indicate boundaries of the nuclei. **(B)** N2a cells stably expressing PrPΔGPI-GFP were treated for 4 h with Apratoxin S9 (100 nM), or Importazole (50 µM), or with both. The cells were fixed and GFP fluorescence was analyzed by SR-SIM. Nuclei were stained with DAPI (Merged). Using FIJI software, the mean nuclear intensity of GFP fluorescence was quantified. The indicated line is the mean of the data set and was analyzed by Kruskal-Wallis test followed by Dunn’s Multiple Comparison Test at 99% confidence interval, *** p=0.0001. **** p=0.00001. At least 50 cells were analyzed per biological replicate (n=3).

### PrP forms partially PK-resistant aggregates in the cytosol but not in the nucleus

To analyze the degradation of PrP in the cytosol and in the nucleus, we treated cells transiently expressing N3PrP with cycloheximide (CHX) to inhibit protein translation. At the beginning of the CHX chase, N3PrP was detected in the nucleus and in the cytosol. After 2 h of CHX treatment, the nuclear fraction of N3PrP had disappeared, whereas N3PrP in the cytosol was still present 6 h after inhibiting protein translation (Fig. 4A). The increased stability may indicate a misfolded conformation of N3PrP in the cytosol. To probe for conformational differences of PrP in the cytosol and nucleus, we performed limited proteolysis experiments. Protein lysates from NLS-PrP-GFP- or NES-PrP-GFP-expressing cells were treated with increasing concentrations of proteinase K (PK) prior to immunoblotting. At PK concentrations that completely digested NLS-PrP-GFP, the cytosolically localized NES-PrP-GFP resisted proteolytic degradation, suggesting that NES-PrP-GFP adopted a more aggregated conformation compared to NLS-PrP-GFP (Fig. 4B). This analysis also revealed that the N-terminal domain of PrP is highly sensitive to proteolytic digestion, consistent with its intrinsically disordered structure.

As another approach to study conformational differences, we recorded fluorescence recovery after photobleaching (FRAP) to quantify protein mobility of NLS-PrP-GFP and NES-PrP-GFP in living cells. Fluorescence recovery of the NES-PrP-GFP assemblies in the cytosol was clearly reduced in comparison to that of the NLS-PrP-GFP assemblies, indicating differences in their material properties (Fig. 4C).

**Figure 4.**
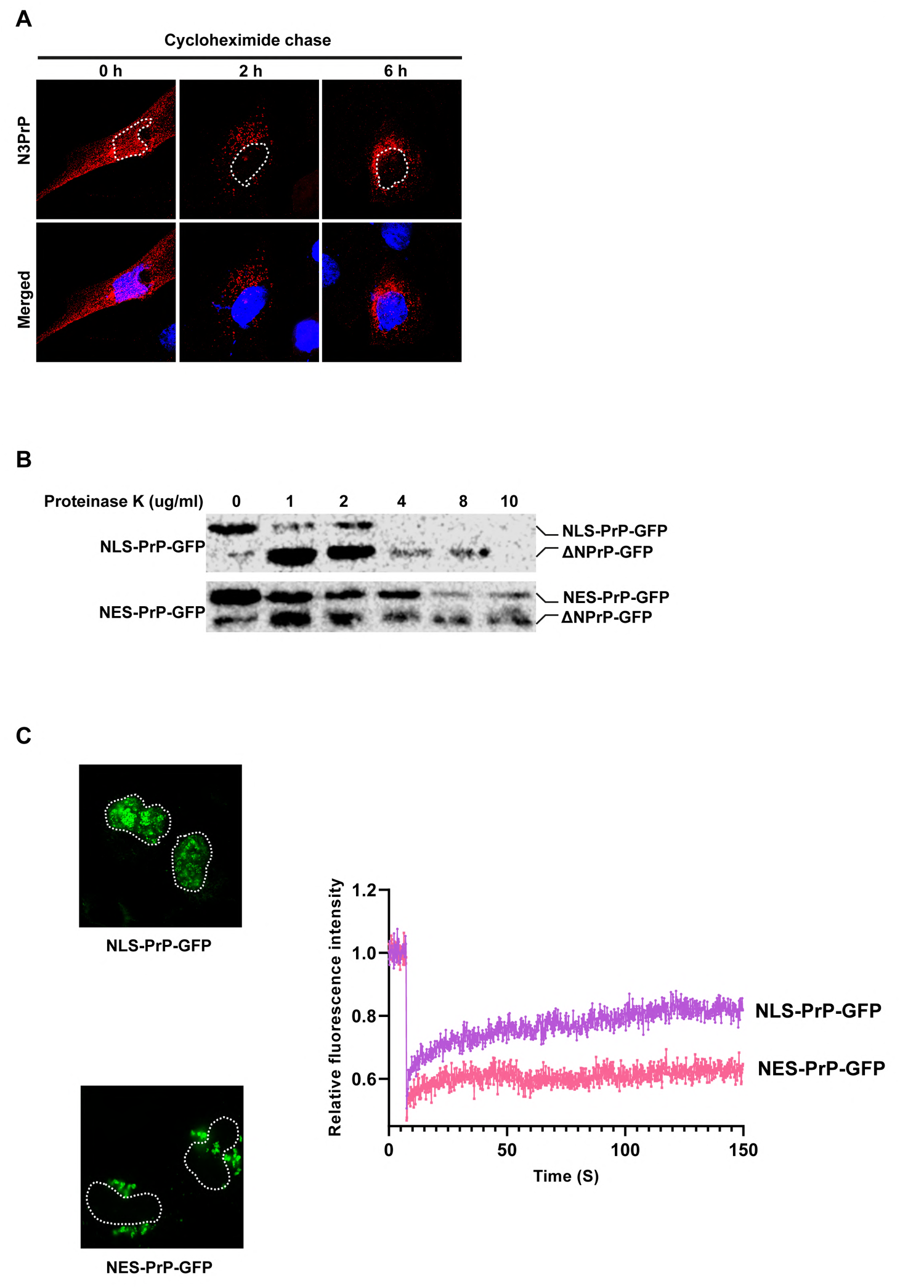
Cytosolic PrP forms partially protease-resistant aggregates. **A. Nuclear PrP is rapidly degraded.** SH-SY5Y cells were transiently transfected with N3PrP. 24 h post-transfection cells were treated for 2 h or 6 h with 50 µg/ml Cycloheximide to inhibit translation. Cells were then fixed, stained with antibodies against PrP and analyzed by SR-SIM. Nuclei were stained with DAPI (Merged). White dotted lines indicate boundary of the nucleus. **B. Cytosolic PrP adopts a partially proteinase-K-resistant conformation.** HEK-293T cells were transfected with NLS-PrP-GFP or NES-PrP-GFP. 24 h post-transfection, cells were lysed and digested with different concentrations of proteinase-K as indicated for 15 minutes at 37°C. Cell lysates were analyzed by immunoblotting using antibodies against PrP. The PrP antibody used (3F4) recognizes the amino acids 106–115, indicating that the faster migrating band represents truncated PrP-GFP molecules devoid of the N-terminally intrinsically disordered domain. **C. Nuclear PrP forms dynamic protein assemblies.** SH-SY5Y cells were transiently transfected with NLS-PrP-GFP or NES-PrP-GFP. 24 h post-transfection, live cells were analyzed by FRAP to probe for the mobility of the PrP molecules. Relative fluorescence intensity was calculated and plotted against the time-period of analysis. Total of 7 cells were analyzed by FRAP for each condition and mean relative fluorescence intensity is shown on the plot. White dotted lines indicate boundaries of the nuclei.

### VCP/p97 prevents aggregation of PrP in the cytosol

Our previous experiments suggested that PrP is inefficiently degraded by proteasomes in the cytosol. We therefore considered the possibility that the interaction with VCP/p97 prevents aggregation of PrP in the cytosol and thereby facilitates importin-β-mediated nuclear import. To study a possible anti-aggregation activity of VCP/p97, we established an *in vitro* aggregation assay with purified components. This assay is based on a recombinant MBP-PrP-GFP fusion protein. The N-terminal MBP (maltose-binding protein) keeps PrP soluble during purification and can be cleaved off by TEV (Tobacco Etch Virus) protease to initiate phase transition of PrP (Fig. 5A) (Kamps et al., 2021). When MBP is cleaved off, PrP-GFP is no longer soluble in physiological buffer (TRIS pH 7.4, 150 mM NaCl) and rapidly aggregates (Fig. 5A, row 1). Aggregation of PrP-GFP after TEV cleavage was prevented in the presence of recombinant VCP/p97 (Fig. 5A, row 2; Fig. EV3A). Upon addition of ATP, PrP-GFP formed aggregates, suggesting that the anti-aggregation activity of VCP/p97 was due to a specific interaction of VCP/p97 with PrP (Fig. 5A, row 2). In line with this notion, the VCP/p97 inhibitor NMS-873, which induced the formation of cytosolic PrP aggregates in cells (Fig. 2A), also induced PrP-GFP aggregation by releasing PrP-GFP from its binding to VCP/p97 (Fig. 5A, row 3).

**Figure 5.**
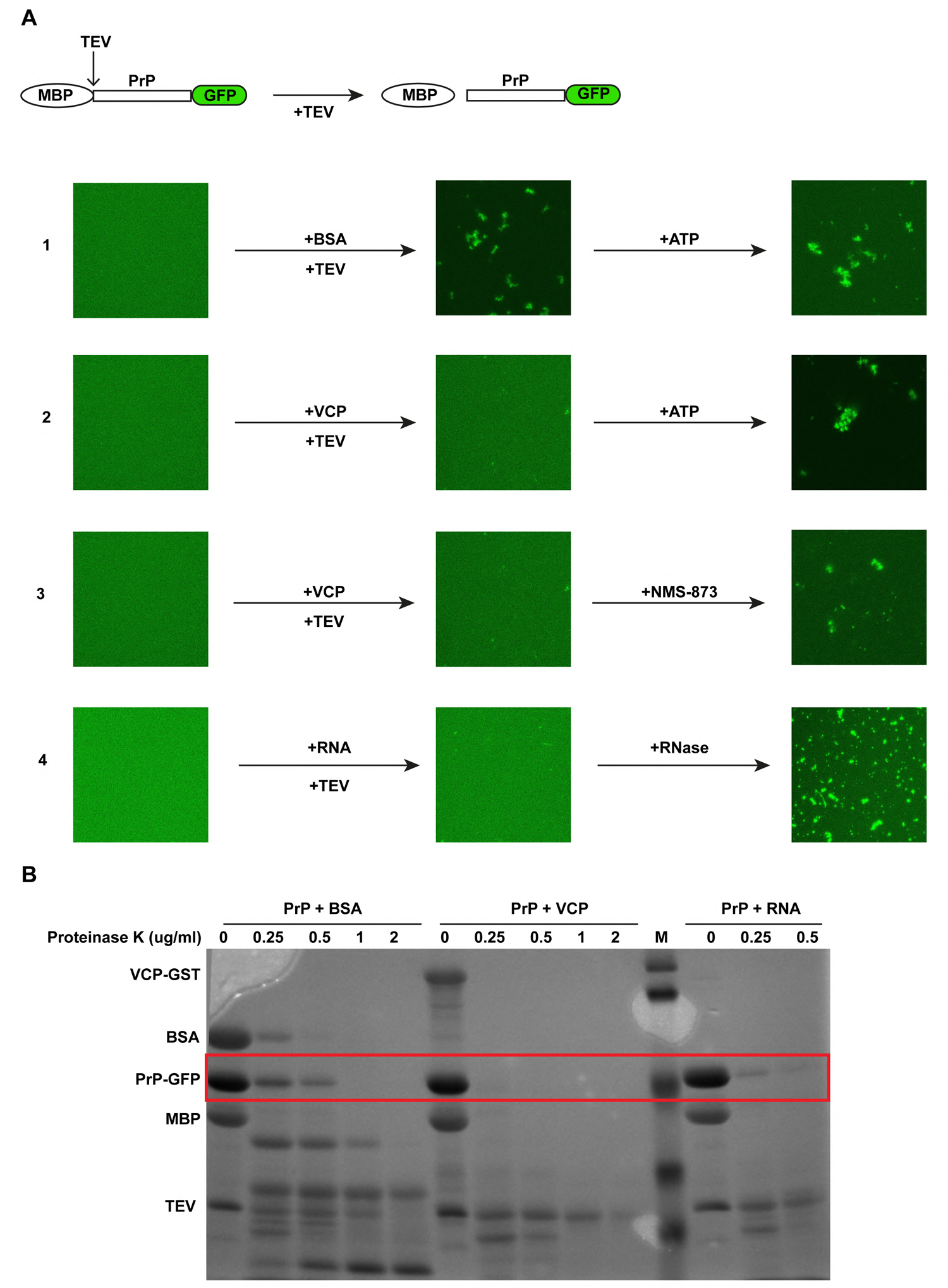
VCP/p97 and RNA prevent a conformation transition of soluble PrP into aggregates. **A. VCP/p97 and RNA inhibit aggregation of PrP-GFP *in vitro***. Scheme of the experimental approach. MBP-PrP-GFP is expressed in *E. coli* and purified as a soluble protein. The N-terminal MBP-tag is removed by TEV protease to induce phase separation (top panel). **1 - 4.** 5 µM MBP-PrP-GFP was incubated in the presence of TEV protease for 1 hour at room temperature in aggregation buffer (Tris pH 7.4, 150 mM NaCl). To analyze aggregation of PrP-GFP fluorescence imaging data were recorded by laser scanning microscopy using Z-stack and processed with Maximum Intensity Projection. **1.** BSA (5 µM) was added together with TEV. After 1 h the reaction was incubated for an additional 1 h in the presence of 2 mM ATP. **2.** Recombinant VCP-GST (5 µM) was added together with TEV. After 1 h the reaction was incubated for an additional 1 h in the presence of 2 mM ATP. **3.** Recombinant VCP/p97-GST (5 µM) was added together with TEV. After 1 h the reaction was incubated for an additional 1 h in the presence of the allosteric VCP/p97 inhibitor NMS-873 (10 µM). **4.** Bulk RNA prepared from HeLa cells was added together with TEV. After 1 h the reaction was incubated for an additional 1 h in the presence of RNase. **B. VCP/p97 and RNA maintain PrP-GFP in a soluble conformation.** 5 µM MBP-PrP-GFP was incubated in the presence of TEV protease for 1 hour at room temperature in aggregation buffer (Tris pH 7.4, 150 mM NaCl) in the presence of either BSA (5 µM), or recombinant VCP-GST5 (5 µM), or bulk RNA (300 ng/µl) prepared from Hela cells. The samples were treated with the indicated concentrations of proteinase-K for 15 minutes at 37°C. The reaction was stopped by addition of 5 mM PMSF and the samples were analyzed by SDS-PAGE and Coomassie brilliant blue staining.

To probe for conformational differences between VCP/p97-bound PrP-GFP and aggregated PrP-GFP, we performed limited proteolysis experiments. In complex with VCP/p97, PrP-GFP was more sensitive to proteolytic digestion compared to aggregated PrP (PrP + BSA), indicating that VCP/p97 acted as a holdase to prevent a conformational transition of soluble PrP-GFP into aggregates (Fig. 5B).

### In the nucleus, RNA prevents phase transition of PrP into aggregates

In the nucleus, NLS-PrP did not interact with VCP/p97 (Fig. 2D), yet, it did not form immobile aggregates, as shown by FRAP recordings (Fig. 4 C). A major difference between the cytosolic and nuclear chemical milieu is the high content of negatively charged RNAs in the nucleus, which can buffer protein aggregation (Maharana et al., 2018). To test for a possible role of RNA in keeping PrP-GFP soluble, we used the *in vitro* aggregation assay described above. Indeed, bulk RNA purified from HeLa cells interfered with the transition of soluble PrP-GFP into aggregates (Fig. 5A, row 4, Fig. EV3A). The anti-aggregation activity was dependent on RNA polymers, since PrP-GFP started to aggregate upon addition of RNase (Fig. 5A, row 4). Moreover, in the presence of RNAs, PrP-GFP was more sensitive to proteolytic digestion, indicating that RNAs acted as negatively charged polymeric cosolute to keep PrP-GFP in a soluble conformation (Fig. 5B).

### The interaction of PrP with VCP/p97 in the cytosol and nuclear targeting is independent of ubiquitination but requires the unstructured N-terminal domain of PrP

Ubiquitination of client proteins has been shown to mediate their interaction with VCP/p97. However, our *in vitro* aggregation assays indicated that VCP/p97 can obviously interact with non-ubiquitinated PrP. To explore this aspect in a cellular context, cells transiently expressing PrPΔGPI-GFP were treated with Apratoxin S9 in combination with an inhibitor of the E1 ubiquitin-activating enzyme (TAK-243). Immunoblotting of whole cell lysates showed that TAK-243 efficiently inhibited ubiquitination of proteins (Fig. 6A). In parallel, we immunoprecipitated TrapVCP and probed the immunopellet for PrP. Notably, non-ER-imported PrPΔGPI-GFP co-purified with TrapVCP also in TAK-243-treated cells, indicating that the PrP-VCP/p97 interaction was independent of ubiquitination (Fig. 6B). Likewise, ubiquitination was not required for nuclear targeting, since PrPΔGPI-GFP localized to the nucleus in cells treated with Apratoxin S9 and TAK-243 (Fig. 6C). This finding raised the question whether proteasomal degradation of PrP in the nucleus is also independent of ubiquitination. We therefore analyzed cells transiently expressing the PrP115X mutant. Non-ER-imported PrP115X is detectable in the nucleus only after proteasomal inhibition (Fig. 1C, 6D). Upon TAK-243 treatment, PrP115X accumulated in the nucleus, similarly to Bortezomib-treated cells, revealing that ubiquitination of PrP is required for its proteasomal degradation in the nucleus.

**Figure 6.**
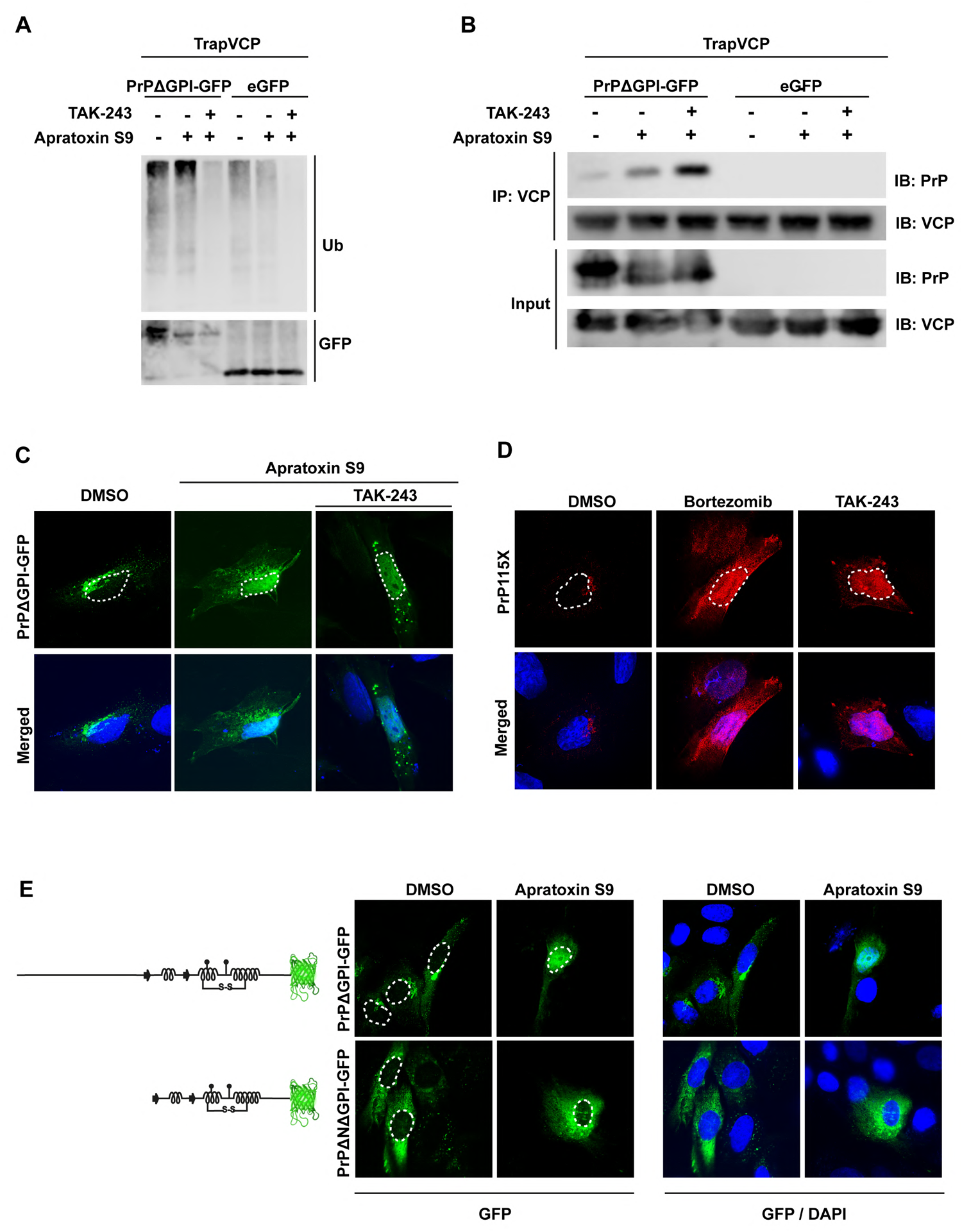
Nuclear targeting of non-imported PrP is independent of ubiquitin but requires the N-terminal unstructured domain of PrP. **A, B. VCP/p97 interacts with non-imported PrP independent of ubiquitination** HEK-293T cells were co-transfected with Strep-tagged E578QVCP (TrapVCP) and PrPΔGPI-GFP, or eGFP. 24 h post-transfection, cells were treated for 3 h with Apratoxin S9 (3 h, 100 nM) and/or TAK-243 (4 h, 1 µM) an inhibitor of ubiquitin activating enzymes (UAE) as indicated. Note that the UAE inhibitor was added 1 h before Apratoxin S9. **(A)** Cell lysates were analyzed by immunoblotting using antibodies against Ubiquitin (P4D1) and GFP. **(B)** Cell lysates subjected to immunoprecipitation under native conditions using Streptactin magnetic beads. Precipitated proteins were then detected by immunoblotting using antibodies against PrP and VCP/p97. **C. Nuclear import of PrP is independent of ubiquitination.** SH-SY5Y cells were transiently transfected with PrPΔGPI-GFP. 24 h post-transfection cells were treated with Apratoxin S9 (3 h, 100 nM), or Apratoxin S9 (3 h, 100 nM) and TAK-243 (4 h, 1 µM). Note that TAK-243 was added 1 h before Apratoxin S9. The cells were fixed and GFP fluorescence was analyzed by SR-SIM. White dotted lines indicate boundaries of the nuclei. Nuclei were stained with DAPI (Merged). **D. Ubiquitination is required for proteasomal degradation of PrP in the nucleus.** SH-SY5Y cells were transiently transfected with PrP115X. 24 h post-transfection cells were treated for 3 h with Bortezomib (1 µM), or TAK-243 (1 µM). Cells were fixed, stained with antibodies against PrP and analyzed by SR-SIM. White dotted lines indicate boundary of the nucleus. Nuclei were stained with DAPI (Merged). **E. The N-terminal intrinsically disordered domain of PrP is required for nuclear import of PrP.** SH-SY5Y cells were transiently transfected with PrPΔGPI-GFP or PrPΔNΔGPI-GFP. 24 h post-transfection cells were treated for 4 h with Apratoxin S9 (100 nM). The cells were fixed and GFP fluorescence was analyzed by SR-SIM. White dotted lines indicate boundaries of the nuclei. Nuclei were stained with DAPI (Merged).

PrP115X, which is efficiently targeted to the nucleus, is composed only of the intrinsically disordered domain of PrP (Fig. 1A, 6D). We therefore wondered whether nuclear targeting of the C-terminal structured part of PrP is dependent on the N-terminal domain. Thus, we analyzed nuclear targeting of PrPΔ23-121 (PrPΔN), which lacks the unstructured N-terminal domain (Gonsberg et al., 2017). In contrast to PrP115X and PrPΔGPI-GFP, PrPΔNΔGPI-GFP, remained in the cytosol after inhibiting ER import and was not translocated into the nucleus (Fig. 6E, EV3B).

As a conclusion, nuclear targeting of non-ER-imported PrP depends on the unstructured N-terminal domain of PrP but not on ubiquitination. For proteasomal degradation of PrP in the nucleus, however, ubiquitination is required.

### Proteotoxic stress in the cytosol disrupts nuclear translocation of non-ER-imported PrP and induces the formation of self-perpetuating PrP aggregates in the cytosol

During our study we noticed that cytosolic PrP aggregates increased over time upon defective ER import, whereas the relative amount of nuclear PrP decreased (Fig. 7A). The cellular localization of N3PrP was analyzed in transiently transfected cells by fluorescence microscopy at 3, 19 and 45 h after transfection. We observed an increase in cytosolic N3PrP aggregates already at 19 h after transfection, whereas the nuclear fraction of N3PrP steadily decreased and was barely detectable at 45 h. This observation led us to hypothesize that the cytosolic PrP aggregates and proteotoxic stress induced by these aggregates disrupt further nuclear import of newly synthesized PrP. To test this possibility experimentally, we induced proteotoxic stress in cells expressing PrPΔGPI-GFP by transiently inhibiting the proteasome with Bortezomib (3 h pretreatment). Then, the reversible proteasome inhibitor was washed out and ER import was inhibited by Apratoxin S9. After additional 4 h the cells were fixed and analyzed by fluorescence microscopy. In cells treated with Apratoxin S9 only, the typical nuclear localization of non-ER-imported PrPΔGPI-GFP was observed. However, a pretreatment with Bortezomib interfered with the transport of PrPΔGPI-GFP into the nucleus and led to the formation of cytosolic PrP aggregates (Fig. 7B). To address possible longer-term effects of a transient proteotoxic stress, we induced the formation of cytosolic PrPΔGPI-GFP aggregates by a short treatment (4 h) with Bortezomib and Apratoxin S9. The cells were then analyzed either directly without recovery or after 48 h recovery in fresh medium (Fig. 7C, D). Remarkably, after 48 h recovery cytosolic PrP aggregates were still detectable. These PrP aggregates obviously recruited non-ER-imported PrPΔGPI-GFP, which was permanently generated even in the absence of Apratoxin S9, indicating a self-perpetuating property that interfered with further nuclear targeting of cytosolic PrP.

**Figure 7.**
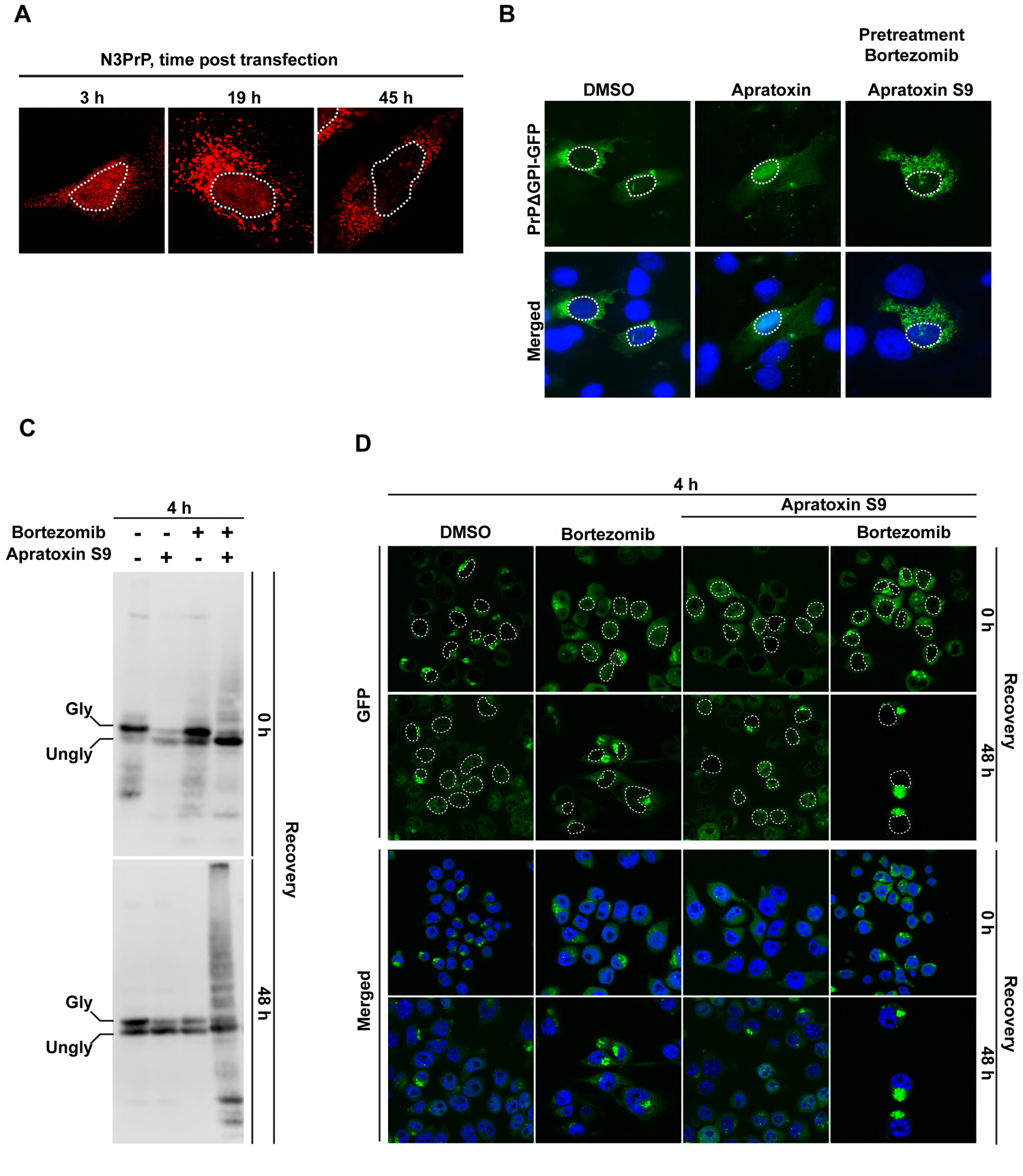
Cytosolic PrP forms self-perpetuating aggregates that disrupt nuclear targeting of PrP. **A. Cytosolic PrP limits the rate of nuclear translocation of newly synthesized PrP.** SH-SY5Y cells were transiently transfected with N3PrP and cultured for the indicated time. Cells were then fixed, stained with antibodies against PrP and analyzed by SR-SIM. Dotted lines represent the nuclear boundary of the analyzed cells. **B. Short-term proteasomal inhibition impairs subsequent nuclear targeting of non-imported PrP.** SH-SY5Y cells were transiently transfected with PrPΔGPI-GFP. 24 h post-transfection cells were pretreated for 3 h with Bortezomib (1 µM). The cells were washed and incubated for 4 h in fresh media containing Apratoxin S9 (100 nM). GFP fluorescence of fixed cells was analyzed by SR-SIM. White dotted lines indicate boundaries of the nuclei. Nuclei were stained with DAPI (Merged). **C, D. Cytosolic PrP forms self-perpetuating aggregates.** **(C, D)** N2a cells stably expressing PrPΔGPI-GFP were plated and treated for 4 h with Apratoxin S9 (100 nM), and/or Bortezomib (0.5 µM) as indicated. **(C)** Cells were lysed either immediately (0 h recovery), or washed with fresh medium and cultured for additional 48 h (48 h recovery). Cell lysates were analyzed by immunoblotting using antibodies against PrP. **(D)** The cells were fixed and GFP fluorescence was analyzed by SR-SIM either directly (0 h recovery), or washed with fresh medium and cultured for additional 48 h (48 h recovery). White dotted lines indicate boundaries of the nuclei. Nuclei were stained with DAPI (Merged).

Interestingly, PrP115X did not form cytosolic aggregates after proteasomal inhibition (Figs. 1C, 6D), suggesting that the C-terminal domain is linked to the formation of cytosolic aggregates. To address this aspect experimentally we analyzed nuclear import of PrP-Y163X, which is linked to inherited prion disease in humans and contains only a part of the C-terminal structured domain (Finckh et al., 2000, Jayadev et al., 2011, Kitamoto et al., 1993, Mead et al., 2013). PrP163X-GFP was imported into the nucleus, however, in contrast to 115X-GFP it also formed cytosolic aggregates after proteasomal inhibition. This finding is emphasizing a critical role of the structured C-terminal domain in driving aggregation of PrP in the cytosol (Fig. EV4).

### Cytosolic but not nuclear PrP causes a proteostasis decline in both the cytosol and nucleus

The FRAP and limited proteolysis experiments revealed that PrP adopts distinct conformations in the cytosol and the nucleus, respectively. We therefore wondered whether these conformational differences translate into specific biological consequences. To monitor the cellular proteostasis capacity, we expressed a mutated version of the conformationally unstable firefly luciferase (Fluc) protein fused to EGFP. Proteotoxic stress leads to misfolding of Fluc-EGFP, resulting in decreased luciferase activity (Blumenstock et al., 2021, Gupta et al., 2011, Park et al., 2013). After co-expression of Fluc-EGFP with either NES- or NLS-PrP the enzymatic activity of Fluc-EGFP was quantified by a luciferase assay. In cells expressing NLS-PrP, luciferase activity was not decreased compared to control cells. In contrast, a significant decline in luciferase activity was observed in cells expressing NES-PrP (Fig. 8A, left panel). To specifically detect compartment-specific alterations in proteostasis, we targeted Fluc-EGFP either to the nucleus (NLS-Fluc-EGFP) or to the cytosol (NES-Fluc-EGFP). The co-expression of NLS-PrP did neither affect the luciferase activity of cytosolic nor nuclear Fluc-EGFP (Fig. 8A). However, PrP accumulation in the cytosol upon NES-PrP expression significantly decreased the luciferase activity of both NLS-Fluc-EGFP and NES-Fluc-EGFP (Fig. 8A, middle and right panel).

**Figure 8.**
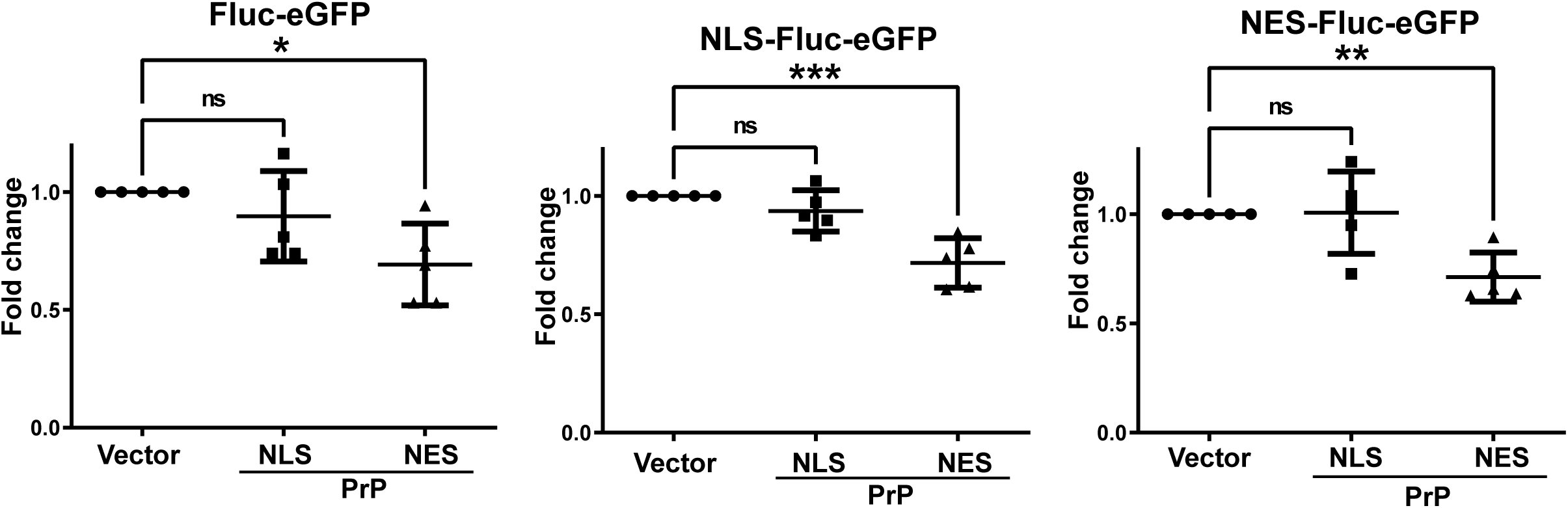
Cytosolic PrP but not nuclear PrP, causes proteostasis decline in the cytosol and nucleus. Fluc-eGFP folding sensors present in both the nucleus and the cytosol (Fluc-eGFP), or only in the nucleus (NLS-Fluc-eGFP), or only in the cytosol (NES-Fluc-eGFP), were coexpressed in HEK cells with NLS-PrPΔGPI, or NES-PrPΔGPI. 24 h post-transfection, cells were lysed, and luminescence in total lysates was measured using a Luminometer. Fold change of luminescence was calculated by standardizing against lysates from cells transfected with an empty vector instead of PrP. The indicated line is the mean of the data set of 5 biological replicates, analyzed with Bonferroni’s Multiple Comparison Test at 95% confidence interval, * p=0.05 ** p=0.005 *** p=0.0005.

## DISCUSSION

Rapid and efficient degradation of non-native protein conformers is a key mechanism in protein quality control to prevent the accumulation of protein aggregates and subsequent proteostasis imbalances. Here we identified a VCP/p97- and importin-dependent quality control pathway that inhibits the formation of toxic cytosolic protein aggregates by targeting non-ER-imported secretory proteins to the nucleus for proteasomal degradation (Fig. 9). By studying PrP as a disease-relevant secretory protein, we found that the intrinsically unstructured N-terminal domain was required for nuclear translocation of non-ER-imported PrP. PrP115X, which comprises only the disordered N-terminal region, was efficiently imported into the nucleus and did not form cytosolic aggregates, not even upon prolonged proteotoxic stress. In contrast, PrPΔ23-121, which consists of the C-terminal structured region, aggregated in the cytosol. Interestingly, the pathogenic C-terminal deletion mutants PrP-Y145X, -Q160X and -Y163X contain a part of the C-terminal structured domain (Finckh et al., 2000, Jayadev et al., 2011, Kitamoto et al., 1993, Mead et al., 2013), whereas PrP deletion mutants that are entirely unstructured, such as PrP-R37X and -Q75X, were found in healthy controls (Minikel et al., 2016). These observations suggest that the presence of the C-terminal structured region impairs nuclear targeting of non-ER-imported PrP and that this correlates with neurotoxic adverse effects. In support of this notion, PrP-Y163X was not efficiently imported into the nucleus and aggregated in the cytosol.

**Figure 9.**
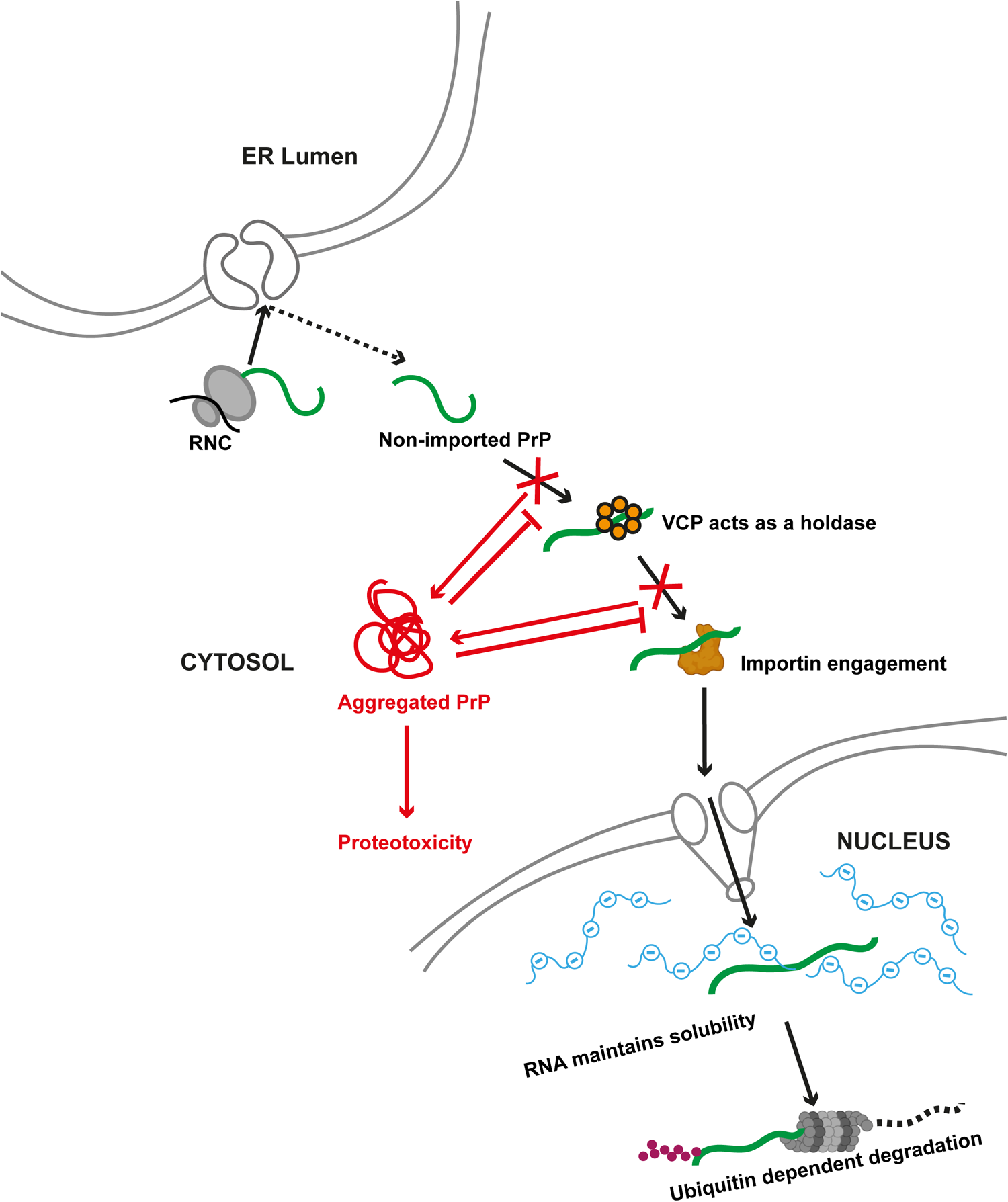
Schematic summary of the findings. Under physiological conditions a sequential interaction of non-ER-imported PrP with VCP/p97 and importins maintain PrP in a soluble conformation in the cytosol and mediates its import into the nucleus. High concentrations of RNA buffer aggregation of PrP in the nuclear compartment and facilitate the ubiquitin-dependent proteasomal degradation of PrP. Under proteotoxic stress conditions the interactions with VCP/p97 and importins are disturbed and PrP forms self-perpetuating cytosolic aggregates that prevent further nuclear targeting of non-ER-imported PrP.

Employing a variety of *in vitro* and *in cellulo* approaches we were able to gain insight into the mechanism of this nucleus-based protein quality control pathway. Firstly, an interaction of PrP with VCP/p97 was required, which did not result in cytosolic degradation of PrP by the proteasome. Instead, VCP/p97 maintained non-ER-imported PrP in a soluble conformation after its release from the ribosome. This activity was independent of VCP/p97 adaptor proteins and PrP ubiquitination. Purified VCP/p97 prevented the aggregation of non-ubiquitinated PrP *in vitro* in the absence of VCP/p97 adaptor proteins. Moreover, an inhibitor of E1 ubiquitin-activating enzymes did not interfere with nuclear targeting of PrP in cellular models. The ATP-induced release of PrP from VCP/p97 facilitated the subsequent interaction of PrP with importin-ß, indicating that VCP/p97 acted as a holdase in this context. Of note, the interaction of PrP with VCP/p97 was a prerequisite for nuclear targeting of PrP by importin-ß. VCP/p97 inhibitors induced aggregation of PrP in the cytosol and prevented its nuclear translocation. The subcellular localization of VCP/p97 in close proximity to ER membranes may allow its rapid binding to PrP after abortive ER import and release from the Sec61 translocon. Our data also add to the notion that importins have a chaperone-like activity (Guo et al., 2018, Hofweber et al., 2018, Qamar et al., 2018), since they not only mediated nuclear translocation of PrP but also prevented PrP aggregation in the cytosol. Surprisingly, non-ER-imported PrP was not efficiently degraded by the proteasome in the cytosol, although it was kept in a soluble conformation by VCP/p97. A possible explanation would be that in the cytosol the VCP/PrP complex is not associated with VCP/p97 cofactors required for targeting clients to the proteasome. Ubiquitination of PrP was dispensable for nuclear translocation of PrP, but required for its proteasomal degradation in the nucleus. What could be the advantage of directing aggregation-prone proteins to the nucleus for proteasomal degradation? One relevant difference between the cytoplasm and nucleoplasm that matters in this context is the increased abundance of mRNAs and noncoding RNAs in the nucleus. Our *in vitro* experiments revealed that RNAs prevented the aggregation of PrP, suggesting that RNAs can act as negatively charged polymeric cosolutes to keep PrP in a soluble conformation.

Earlier work from the Hegde lab revealed that even under physiological conditions, 10 to 20% of newly synthesized PrP are not imported into the ER and rapidly aggregate in the cytosol after proteasomal inhibition (Rane et al., 2004). This fraction is obviously efficiently cleared by the nuclear-based quality control pathway, highlighting its important function in maintaining proteostasis under physiological conditions. However, previous studies did not consider the fact that PrP aggregates formed in the cytosol upon proteasomal inhibition occur as a consequence of inefficient nuclear import. Moreover, we observed that PrP aggregates once formed in the cytosol self-perpetuate and are no longer translocated into the nucleus, even when proteasomal function is restored. Our data suggest that impaired nuclear translocation of cytosolic PrP and its aggregation in the cytosol PrP are pathophysiologically relevant. Cytosolic and nuclear PrP assemblies were different regarding their material properties and proteotoxic potential. Limited proteolysis and FRAP experiments revealed that nuclear PrP was in a soluble conformation and mobile, whereas cytosolic PrP formed partially PK-resistant, immobile aggregates. Notably, only cytosolic but not nuclear PrP assemblies caused an imbalance in proteostasis.

In a wider perspective, our study underscores the concept that mislocalization of aggregation-prone proteins can trigger the accumulation of aggregates with adverse effects on the proteostasis balance (Hartl, 2017, Juszkiewicz & Hegde, 2018, Yanagitani et al., 2017). It is quite remarkable that cells have the capacity to respond to such a mislocalization with another “mislocalization” that cannot restore the function of the missorted or orphaned proteins but at least prevent a toxic gain of function. Redirecting non-ER-imported misfolding-prone proteins to the nucleus, where they can be degraded more efficiently, counteracts the accumulation of harmful species in the cytosol, which seems to be particularly vulnerable to proteostasis dysregulation. Strikingly, the nucleus has recently been identified as a destination for the proteasomal degradation of non-imported mitochondrial precursors (Shakya et al., 2021), suggesting that nuclear redirection is a more general quality control pathway for proteins mislocalized in the cytosol. It will now be interesting to characterize both shared and specific key players in the nucleus-associated degradation pathway of proteins upon defective import into the ER or mitochondria.

## Materials and Methods

### DNA constructs

Plasmid maintenance and amplification was carried out using *Escherichia coli* TOP10^©^ (ThermoFisher Scientific). All PrP constructs were generated by standard PCR cloning techniques and are based on the coding region of mouse PRNP (GenBank accession number M18070) modified to express PrP-L108M/V111M, allowing detection by the monoclonal antibody 3F4. All the mammalian expression plasmids were cloned into pcDNA3.1(+)-Neo vector (Invitrogen^TM^ V79020). PrPΔGPIGFP: aa 1-226 tagged with GFP at the C-terminus; PrPΔGPI: aa 1-226; NES-PrP-GFP: aa 1-226 tagged with GFP at the C-terminus and with a nuclear export signal at the N-terminus; NLS-PrP-GFP: aa 1-226 tagged with GFP at the C-terminus and with a nuclear localisation signal at the N-terminus; NES-PrP: aa 1-226 tagged with a nuclear export signal at the N-terminus; NLS-PrP: aa 1-226 tagged with a nuclear localisation signal at the N-terminus; N3PrP: mouse PrP with a mutated ER signal peptide (Kim et al., 2002); PrP115X: aa 1-114; PrPΔNΔGPIGFP: aa121-226 tagged with GFP at the C-terminus; ΔSPPrPΔGPIGFP: aa 23-226 tagged with GFP at the C-terminus; were cloned into pcDNA3.1(+)-Neo via HindIII and XhoI. The modified signalling sequences are as follows: NLS: 5’ ATGCCACCAAAAAAAAAAAGAAAAGTT 3’ NES: 5’ ATGCTGGAACTTCTGGAGGATTTGACACTG 3’ Generation of pMAL-MBP-TEV-PrP-eGFP-TEV-His_6_ is described elsewhere(Kamps et al., 2021). pcDNA5FRT/TO-p97-EQ-mycStrep, pcDNA5FRT/TO-p97wt-mycStrep, and pGEX-6P-1-p97 were kind gifts from Hemmo Meyer (Ritz et al., 2011). FlucGFP, NLS-FlucGFP, and NES-FlucGFP were kind gifts from F. Ulrich Hartl and Irena Dudanova (Blumenstock et al., 2021, Gupta et al., 2011, Park et al., 2013)

### Cell culture and transfection

#### HEK293T, HeLa cells, mouse embryonic fibroblasts (MEFs)

Cells were cultured in Dulbecco’s modified Eagle’s medium (DMEM) supplemented with 10%(v/v) foetal bovine serum (FBS) and 100 IU/ml penicillin and 100 µg/ml streptomycin sulphate.

#### Human neuroblastoma SH-SY5Y cells

cultured in Dulbecco’s modified Eagle’s medium F-12 (DMEM/F12) supplemented with 15% (v/v) foetal bovine serum (FBS), 100 IU/ml penicillin, 100 µg/ml streptomycin sulphate and 1 x MEM non-essential amino acids solution (Gibco TM).

#### Mouse neuroblastoma N2a Cells

Cells were cultured in Minimum Essential Medium (MEM) supplemented with 10%(v/v) foetal bovine serum (FBS) and 100 IU/ml penicillin and 100 µg/ml streptomycin sulphate. N2a stable cell lines were maintained in 0.1% Puromycin (Santa Cruz, Germany). N2a stably expressing PrPΔGPIGFP were established by using pHulk plasmid system (Haase et al., 2010).

All cell lines were grown in humidified conditions at 37°C with 5% CO2 and passaged when cells reached 80% confluence. For inhibitor treatment, cells were incubated in complete growth medium with the indicated drug at 37°C.

Unless described otherwise, transfections were performed using the following procedures: For SH-SY5Y and MEF cells Lipofectamine and Plus Reagent (Invitrogen) and for HeLa cells Lipofectamine 2000 (Invitrogen) was used according to the manufacturer’s instructions. HEK293T cells were transfected using PEI (polyethylenimine). DNA and PEI were mixed in a 1:4 ratio (µg:µl) in Opti-MEM and incubated for 15 min at room temperature. The DNA-PEI mixture was then added to the cells.

### Primary neuron culture

Primary neurons were prepared following the protocol from Life Technologies, with modification. All media and solutions were sterile filtered using 0.22 µm membrane filter. Cortices were explanted from embryos 16-18 DPC (day post coitum), carefully washed thrice with ice cold Dissection Buffer (9.9 mM HEPES, 137 mM sodium chloride, 5.4 mM potassium chloride, 0.17 mM sodium phosphate dibasic anhydrous, 0.22 mM potassium phosphate monobasic anhydrous, 33.3 mM D-glucose, 43.8 mM sucrose, pH 7.4) and treated with 1 mg acetyl trypsin for 10 min at 37°C in the water bath. Cortices were then rinsed three times in Dissection Buffer, and 100 U DNAse-I was added. Complete dissociation was performed manually using a series of increasingly smaller pipette tips, followed by straining through a 70 µm filter (EASYSTRAINER, Greiner Bio One, Essen, Germany) to remove larger tissue residues and diluted with neurobasal medium (with 10% FBS, and 0.5 mM glutamine). The number of cells was determined using a haemocytometer and the neurons were plated on poly L-lysine coated plates at densities 1.3× 10^4^ and 1.3× 10^6^ cells/well in 24 well plates and 60 mm dishes, respectively. The medium was changed to neurobasal medium supplemented with B-27 supplement and 0.5 mM glutamine after 50 mins. The primary cortical neurons were cultured in a humidified incubator at 37 °C with 5% CO_2._

5 DIV (day in-vitro) neurons were transfected using Lipofectamine 2000 following the manufacturer’s protocol and analysed for immunofluorescence after 48 h.

### Immunofluorescence and confocal microscopy

Transfected cells were washed with phosphate buffered saline (PBS, Gibco, UK) and then fixed using 4% (w/v) paraformaldehyde for 10 mins. Cells were permeabilized using 0.2% triton X-100 in PBS for 10 mins and washed with PBS. For blocking, cells were incubated for 1 hour in 5% normal goat serum in PBS and incubated in 0.1-0.2% (v/v) primary antibody in the blocking solution overnight at 4°C. Cells were then washed with PBS and incubated in the corresponding 0.1% (v/v) secondary antibody in PBS for 1 hour at 25°C. The secondary antibody was removed and washed away using PBS and 0.2% Tween-20 in PBS for 5 mins. All coverslips were incubated in 0.1% (v/v) DAPI in Millipore, washed with Millipore water and then mounted using Fluoromount-G^TM^ (Invitrogen, USA). For non-permeabilized cells, after fixation, coverslips were counterstained with DAPI and mounted.

Fluorescence images were acquired using the ELYRA PS.1 microscope equipped with an LSM880 (Carl Zeiss, Jena) and a 20x, 63x oil or 100x oil immersion objective. Super-resolution images were generated by structured illumination microscopy (SR-SIM) using 405, 488 and 561 nm widefield laser illumination. SIM confocal images were processed using the ZEN2.3 software (Carl Zeiss, Jena). For the quantification of nuclear fluorescence intensities, confocal images were acquired with the imaging settings kept uniform among replicates.

Fiji (https://imagej.net/software/fiji/) was used to measure the nuclear fluorescence intensity by using DAPI as a marker for nuclear mask and custom macros written for automated image analysis which can be made available by request. The *in vitro* samples were analysed as described previously (Ahlers et al., 2021, Kamps et al., 2021).

### Immunoblotting

Proteins were fractionated by SDS-PAGE and transferred to nitrocellulose or polyvinylidene difluoride membranes by electroblotting. The nitrocellulose membranes were blocked with 5% non-fat dry milk or 5% BSA in TBST (TBS containing 0.1% Tween 20) for 60 min at room temperature and subsequently incubated with the primary antibody diluted in blocking buffer for 16 h at 4°C. After extensive washing with TBST, the membranes were incubated with horseradish peroxidase-conjugated secondary antibody for 60 min at room temperature. Following washing with TBST, the antigen was detected with the enhanced chemiluminescence (ECL) detection system (Promega) as specified by the manufacturer with Azure Sapphire Biomolecular Imager (Azure Biosystems, USA).

### Biochemical analyses

#### Preparation of whole cell lysates and nuclear/cytosolic fractions

To prepare whole cell lysates, the cells were harvested, centrifuged at 1000xg for 5 mins, washed with PBS and lysed in 2X Laemmli sample buffer. Meanwhile, nuclear/cytosolic fractions were obtained using a nuclear extraction kit (Millipore, 2900). All samples were boiled for 5 mins prior to analysis.

#### PNGase-F digestion and glycosylation assessment

In a 6-well plate, HEK293T cells were seeded, grown overnight, and transfected with PrPΔGPI-GFP. After 16 h, the cells were treated with or without 100 nM Apratoxin S9 for 4 h. The cells were harvested (1000xg, 5 mins), lysed in 1% Triton-X100 in PBS, vortexed and incubated on ice for 10 mins. The lysates were centrifuged at 18000xg for 10 mins and the resulting supernatants were treated with PNGase-F (New England Biolabs^inc^) for 3 h at 37°C. The samples were analyzed through sodium dodecyl sulfate polyacrylamide gel electrophoresis (SDS PAGE) using a 10% SDS PAGE gel, and immunoblotted by using an antibody, 3F4, against PrP.

#### Co-immunoprecipitation

In 6-cm dishes, HEK293T cells were seeded and co-transfected with PrPΔGPI and p97Wt-mycStrep or p97-EQ-mycStrep. 24 h post-transfection, cells were harvested (1000xg, 5 mins), lysed with 500 µl lysis buffer (150 mM KCl, 5 mM MgCl2, 50 mM Tris-HCl pH 7.4, 1% Triton X-100, 5% glycerol, 2 mM β-mercaptoethanol supplemented with cOmplete EDTA-free protease inhibitors and PhosSTOP, Roche, Basel, Switzerland), gently resuspended by pipetting and incubated on ice for 20 mins. The lysates were centrifuged at 18000xg for 15 mins at 4°C and the supernatant (input) was collected in separate tubes. MagStrep “type3” XT beads (IBA Lifesciences GmbH, Göttingen) pre-washed with lysis buffer were added to the supernatants and incubated for 2 hours at 4°C with gentle rotation. The flowthrough was removed, and the beads were washed thrice with 500 µl lysis buffer. Finally, the beads were boiled with 50 µl 2x Laemmli sample buffer for 5 mins to elute the immunocomplex. All samples were fractionated by SDS PAGE gels and further analysed through immunoblotting against VCP and PrP.

#### Luciferase assay

HEK293T cells were seeded and co-transfected with Fluc-GFP(Wt) / NLSFluc-GFP(Wt) / NESFluc-GFP(Wt) and NLS-PrPΔGPI / NES-PrPΔGPI / pcDNA3.1(+) empty vector control. 24 h post-transfection, cells were harvested (1000xg, 5 mins), lysed in 100 µl 1X reporter lysis buffer (Promega, USA), vortexed and centrifuged (13000 rpm, 4°C, 15 mins). The resulting supernatant (20 µl) was added into a 96-well plate in quadruplicates. Before loading the plate into the Cytation5 reader (BioTek, Germany), the injectors were washed following the manufacturer’s protocol and primed with the luciferase assay substrate (Promega, USA) diluted in water (1:10). 100 µl of the substrate was injected per well and the luminescence was measured at 557 nm. The empty vector controls were used to normalize all the samples within the same transfection group and plotted to calculate the fold change in luminescence of the Fluc reporter.

#### Fluorescence after photo-bleaching (FRAP) analyses

SH-SY5Y cells were seeded on a 35 mm IBIDI µ-Dish and transfected with NLS-PrP-GFP or NES-PrP-GFP. 24 h post-transfection, cells were imaged in Invitrogen™ Live Cell Imaging Solution. ZEN2.1 bleaching and region software module and Plan-Apochromat 100× numerical aperture 1.46 oil differential interference contrast M27 objective was used for imaging the cells. For each cell analysed, two circular regions of interest were chosen. One region was bleached with 100% laser power and a pixel dwell time of 8.71 ms, with a scan time of 111.29 ms and the other region was used as the reference signal. Relative fluorescence intensity (RFI) was calculated at time t was calculated using following equation RFI=I_BL_(t)/I′_BL_ **/** I_Ref_(t)/I′_Ref_ where I_BL_(t) and I_Ref_(t) are the intensities measured at time t in the photobleached region and the reference region, respectively. I′_BL_ and I′_Ref_ are the intensities measured before photobleaching.

#### Proteinase K digestion

HEK293T cells were seeded and transfected with NLS-PrP-GFP or NES-PrP-GFP. 24 h post-transfection cells were lysed in 200 µl lysis buffer (1% Triton-X100 in 50mM HEPES pH 7.4, 150mM NaCl) on ice for 15 mins. The lysate was cleared at 17000xg for 10 mins and equal amounts of the supernatant (20 µl) were digested with the indicated concentrations of Proteinase-K for 15 mins at 25°C. The reaction was stopped with 5mM PMSF (phenylmethylsulfonyl fluoride), mixed with Laemmli sample buffer and boiled for 1 min. Samples were analysed by immunoblotting against a PrP antibody specific to C-terminus (POM-15) and GFP.

#### Expression and purification of recombinant proteins

Plasmid maintenance, bacterial expression, and purification of MBP-PrP-GFP was performed as described earlier (Kamps et al., 2021). For expression and purification of GST-tagged VCP, transformed E. coli BL21 (DE3) were grown to an OD 600 of 0.6–0.8 before induction with 0.5 mM isopropyl-β-d-thiogalactopyranoside and a change in temperature to 18 °C overnight. Cells were harvested and resuspended in lysis buffer (50 mM HEPES pH 8.0, 150 mM KCl, 2 mM MgCl_2_, 5% glycerol) with 20 ml/l of culture volume, protease inhibitor cocktail and 1 mg/ml of Lysozyme is added to the suspension and stirred with a stir-bar gently for 30 mins at 4°C and lysed by a French press. After centrifugation for 60 min at 20,000 g at 4°C, the lysate was cleared with a 0.8µm filter. GST tagged VCP/p97 was affinity-purified with a GSTTrap FF column (GE Healthcare) with a flow rate of 1-5 ml/min on an ÄKTA purification system. The column was washed with the lysis buffer and the bound proteins were eluted using the lysis buffer but with 20 mM glutathione. Adding glutathione changes the pH, buffer needs to be adjusted to pH7.4 – 8.0 before use. Eluted fractions were assessed for protein content by SDS-PAGE. Desired fractions were pooled, concentrated to <5 ml for injection from loop onto gel filtration column, and further purified using HiLoad 16/600 Superdex 200 pg at 1 ml/min flowrate with gel filtration buffer (50 mM HEPES pH 7.4, 150 mM KCl, 2 mM MgCl_2_, 5% glycerol, 1 mM DTT). Eluted fractions were checked for protein content by SDS-PAGE. Desired fractions were pooled, concentrated to the desired concentration, flash frozen in liquid nitrogen, aliquoted and stored at −80°C. The protein concentration was determined using the absorbance at 280 nm and the extinction coefficient of each protein by Nanodrop 2000 (Thermo Fisher).

#### Sample preparation for *in vitro* aggregation assay

Protein samples were thawed and centrifuged (20000 g, 10 min, 4 °C) to remove aggregates. Afterwards, the buffer was exchanged to 10 mM Tris pH 7.4 using Vivaspin 500 columns with 30000 Dalton cut off (Sartorius Stedim biotech). The samples were centrifuged (12000 g, 7 min, 4 °C) five times for complete buffer exchange and then the protein concentration was determined by NanoDrop 2000. To induce aggregation of PrP-GFP the buffer was supplemented with 150 mM NaCl and incubated with TEV protease for 1 h before microscopy.

#### Limited proteolysis of recombinant PrP-GFP

5 µM MBP-PrP-GFP in aggregation buffer was cleaved with TEV protease in the presence of BSA, GST-VCP or bulk RNA prepared from HeLa cells. The samples were treated for 15 min at 25°C with increasing concentrations of proteinase K as indicated. After stopping the reactions with 5 mM PMSF (phenylmethylsulfonyl fluoride), the samples were analyzed by SDS-PAGE and Coomassie brilliant blue staining.

#### Statistical analysis

Statistical analyses for the luciferase assays, and nuclear GFP intensity quantification were performed using one-way ANOVA (Kruskal-Wallis test with Dunn’s post-test at 95% confidence interval) and t-test (Mann-Whitney test, two-tailed at 95% confidence interval), respectively using the GraphPad PRISM 9.5 software.

## ACKNOWLEDGEMENTS

We thank Nuwin Mohamad for preparing primary cortical neurons, Maike Kumper for establishing the N2a stable cells line expressing PrPΔGPI-GFP, Hemmo Meyer for the VCP/p97 expression plasmids and helpful discussions, Matthias Kracht for the protocol for the expression and purification of recombinant GST-VCP/p97, and F. Ulrich Hartl and Irena Dudanova for the Fluc-eGFP plasmids.

This work was funded by the Deutsche Forschungsgemeinschaft (DFG, German Research Foundation) under Germany’s Excellence Strategy – EXC 2033 – 390677874 – RESOLV (to JK, JT and KFW) and TA 167/6-3 (to JT). Chemical synthesis was supported by the National Institutes of Health, RM1GM145426 and the Debbie and Sylvia DeSantis Chair professorship (to HL).

## CONFLICT OF INTEREST

H. Luesch is co-founder of Oceanyx Pharmaceuticals, Inc., which has licensed patents and patent applications related to apratoxins. The other authors declare that they have no conflict of interest.

---

**Figure EV1.**
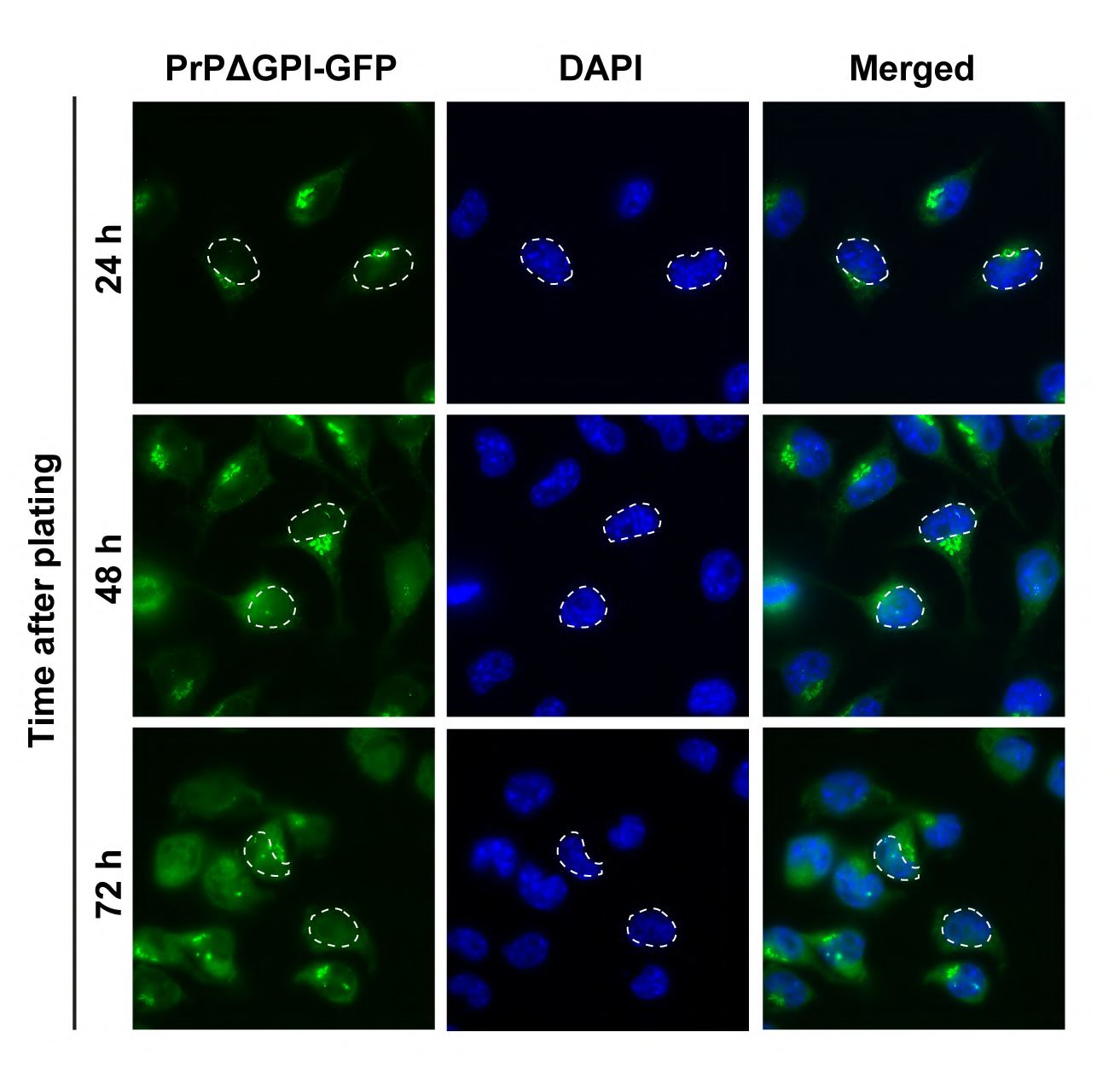
Nuclear targeting of non-imported PrP under physiological conditions in stably transfected cells. N2a cells stably expressing PrPΔGPI-GFP were plated and cultured in complete medium for the indicated time. The cells were fixed and GFP fluorescence was analyzed by SR-SIM. White dotted lines indicate boundaries of the nuclei. Nuclei were stained with DAPI (Merged).

**Figure EV2.**
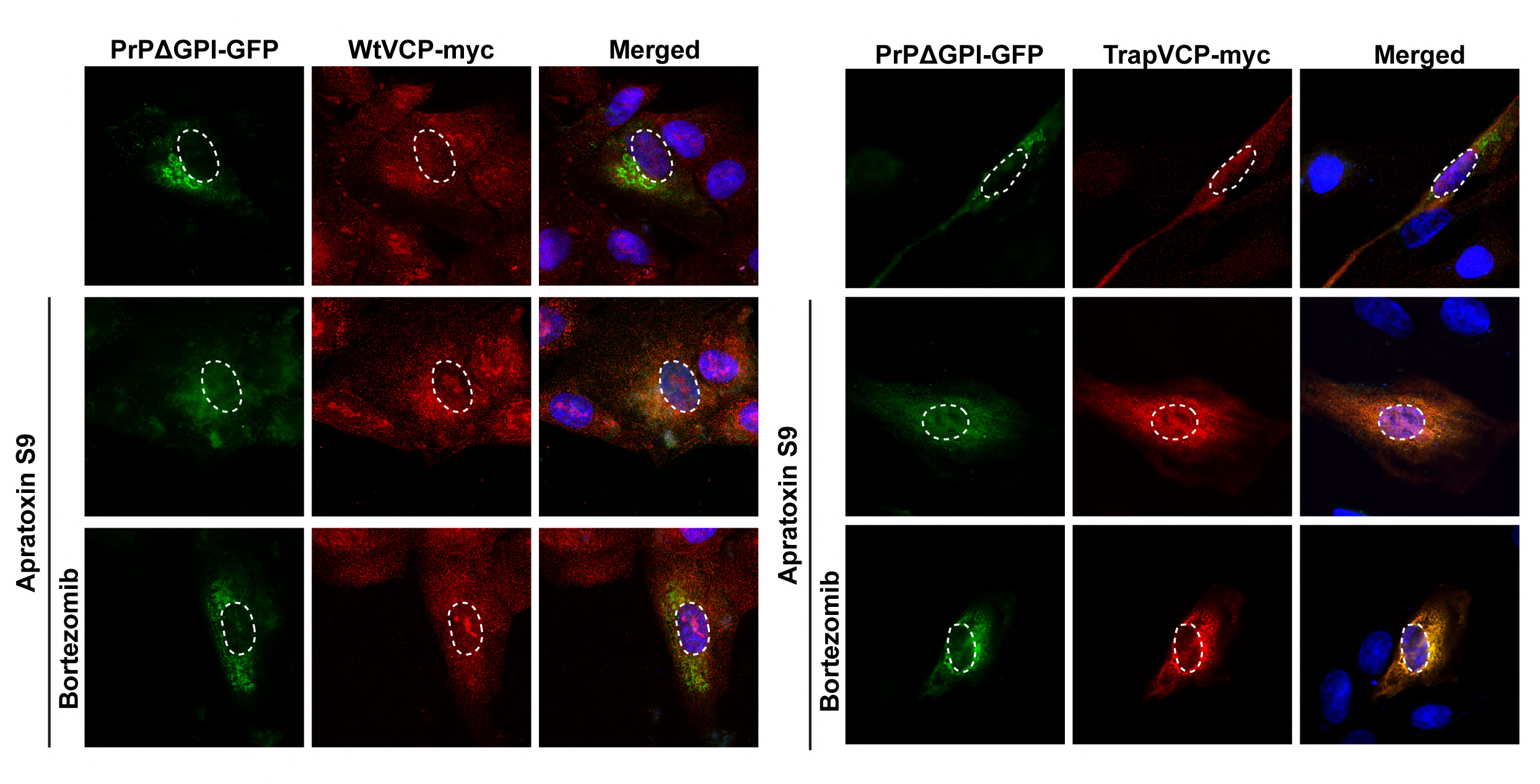
VCP/p97 is located in the cytosol and the nucleus. SH-SY5Y cells were transiently co-transfected with PrPΔGPI-GFP and wild type VCP-myc or TrapVCP-myc. 24 h post-transfection cells were treated for 4 h with Apratoxin S9 (100 nM, 4 h), or Apratoxin S9 and Bortezomib (1 µM). Cells were fixed, stained with antibodies against myc and analyzed by SR-SIM. White dotted lines indicate boundaries of the nuclei. Nuclei were stained with DAPI (Merged).

**Figure EV3.**
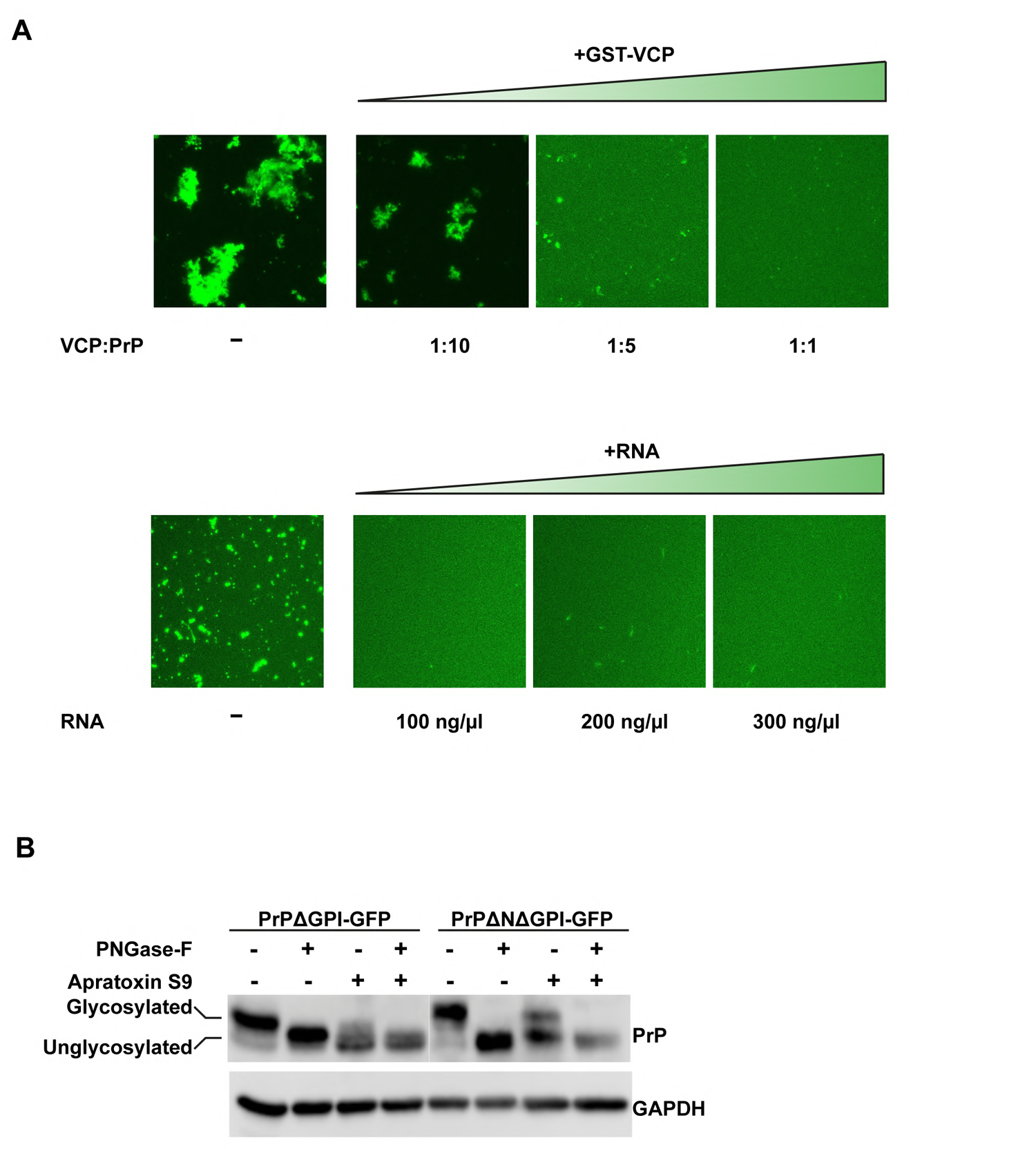
**A. VCP/p97 and RNA inhibit aggregation of PrP-GFP *in vitro.*** 10 µM MBP-PrP-GFP was incubated in the presence of TEV protease for 1 hour at room temperature in aggregation buffer (Tris pH 7.4, 150 mM NaCl). To analyze aggregation of PrP-GFP fluorescence imaging data were recorded by laser scanning microscopy using Z-stack and processed with Maximum Intensity Projection. **Upper row:** increasing amounts of VCP-GST were added together with TEV. **Lower row:** increasing amounts of bulk RNA prepared from HeLa cells were added together with TEV **B. Apratoxin S9 interferes with ER import of PrPΔNΔGPI-GFP.** HEK-293T cells were transiently transfected with PrPΔGPI-GFP or PrPΔNΔGPI-GFP. 24 h post-transfection cells were treated for 4 h with Apratoxin S9 (100 nM). Cells were lysed, digested with PNGase-F, or left untreated and analyzed by immunoblotting using antibodies against PrP and GAPDH (loading control).

**Figure EV4.**
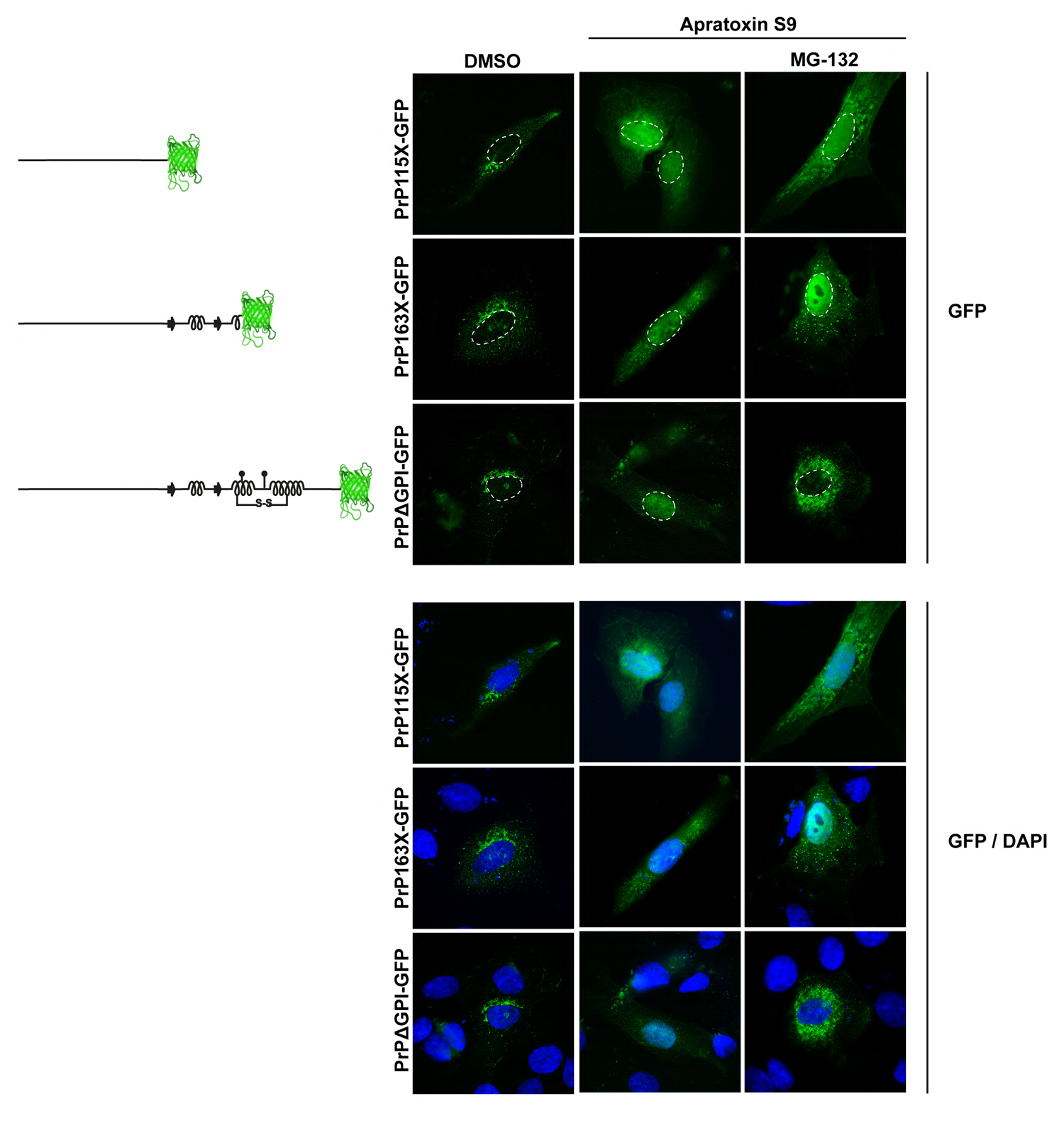
**The C-terminal structured domain of PrP drives aggregation in the cytosol** SH-SY5Y cells were transiently transfected with PrP115X-GFP, or PrP163X-GFP, or PrPΔGPI-GFP. 24 h post-transfection cells were treated for 4 h with Apratoxin S9 (100 nM) and MG-132 (30 µM). The cells were fixed and GFP fluorescence was analyzed by SR-SIM. White dotted lines indicate boundaries of the nuclei. Nuclei were stained with DAPI.

